# Design principles of acidic transcriptional activation domains

**DOI:** 10.1101/2020.10.28.359026

**Authors:** Max V. Staller, Eddie Ramirez, Alex S. Holehouse, Rohit V. Pappu, Barak A. Cohen

## Abstract

Acidic activation domains are intrinsically disordered regions of transcription factors that bind coactivators. The intrinsic disorder and low conservation of activation domains have made it difficult to identify general rules governing their function. To address this problem, we designed thousands of variants in seven acidic activation domains and measured their activities with a new high-throughput assay in human cell culture. From these data we identify three design principles of acidic activation domains: hydrophobic motifs must be balanced by acidic residues; acidic residues make large contributions to activity when they are adjacent to motifs; and motif composition reflects the structural constraints of coactivator interactions. These design principles motivated a simple predictor of activation domains in the human proteome: scanning for clusters of aromatic and leucine residues embedded in regions of high acidity identifies known activation domains and accurately predicts new ones. Our results support a flexible model in which acidic residues solubilize hydrophobic motifs so that they can interact with coactivators.

## Introduction

Transcription factors (TFs) activate gene expression with a DNA binding Domain (DBD) and an activation domain (AD). DBDs are structured, evolutionarily conserved and bind related DNA sequences (Latchman, 2008). In contrast, ADs are intrinsically disordered regions (IDRs), poorly conserved, bind structurally diverse coactivator subunits and frequently undergo coupled folding and binding (Dyson and Wright, 2016). ADs can form clusters in the nucleus, which have been called hubs or condensates (Boija et al., 2018; Chong et al., 2018), but this clustering is separable from AD activity (Powers et al., 2019; Trojanowski et al., 2021). Bioinformatics tools can predict DBDs from protein sequence, but there are no analogous tools for predicting ADs (El-Gebali et al., 2019; Finn et al., 2016). Consequently, when we sequence a new genome we can predict candidate TFs by scanning for DBDs, but we cannot predict which candidate TFs are activators.

Although high acidity (net negative charge) is a common feature of many ADs, acidity is not sufficient for accurate prediction and the literature is conflicted on the functional role of the acidic residues. The first identified ADs contained many acidic residues, leading Paul Sigler to propose the “Acid Blob and Negative Noodle” model (Hope and Struhl, 1986; Ma and Ptashne, 1987; Sigler, 1988). However, site directed mutagenesis soon showed that hydrophobic residues made larger contributions to activity than acidic residues (Cress and Triezenberg, 1991), and in some cases all acidic residues can be removed without loss of activity (Brzovic et al., 2011; Drysdale et al., 1995). While the combination of acidity and hydrophobicity remains an attractive feature for predicting ADs, the leading motif model has low specificity and low sensitivity (Piskacek et al., 2007). Two recent attempts to use deep learning neural networks in yeast, one on random sequences (Erijman et al., 2020) and the other on extant ADs (Sanborn et al., 2020), perform well by cross validation metrics, can identify known ADs and predict new ones. However, it remains challenging to extract mechanistic insights from these “black box” models. Here, we take a complementary approach: we tested a mechanistic model and used our results to build an accurate predictor.

Based on our work in yeast (Staller et al., 2018), we developed an *Acidic Exposure Model* for AD function: acidity and intrinsic disorder keep hydrophobic motifs exposed to solvent where they are available to bind coactivators (Figure 1A). Left to their own devices, hydrophobic residues will interact with each other and drive intramolecular chain collapse, suppressing interactions with coactivators. Surrounding the motifs with acidic residues that repel one another exposes the hydrophobic residues to solvent, promoting interactions with coactivators. We developed this Acidic Exposure Model in yeast and here, test predictions of this model in human cell culture.

**Figure 1.**
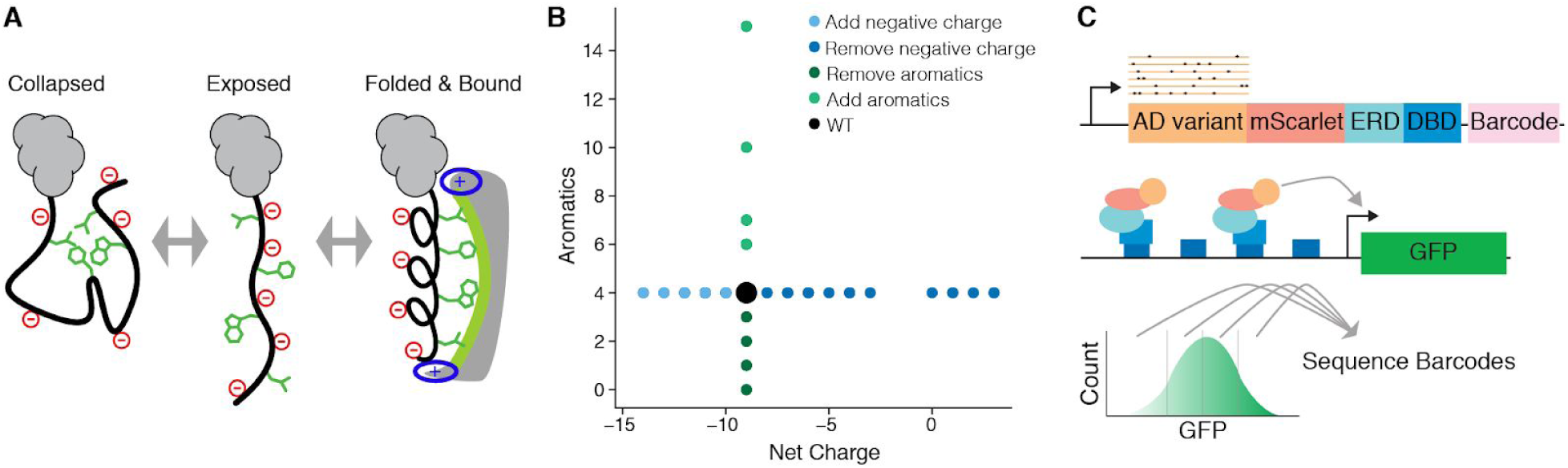
A high throughput assay for measuring the activities of AD variants in parallel. A) In the Acidic Exposure Model, ADs fluctuate between collapsed and exposed states. Exposed ADs can bind coactivators and partially fold. B) Rationally designed mutations add and remove aromatic residues or vary net charge. C) The high-throughput AD assay uses a synthetic DNA binding domain (DBD), an estrogen response domain (ERD), a GFP reporter, FACS and barcode sequencing.

We used a new high-throughput reporter system to test more than 3500 variants in seven known ADs. We designed these variants to interrogate specific aspects of the Acidic Exposure Model. Our results reveal the ways in which acidic and hydrophobic residues are balanced within ADs. Based on these results, we found that scanning the proteome for clusters of eight amino acids (acidic, basic, aromatic and leucine residues) was sufficient to accurately predict new and known ADs. We synthesize our findings in three design principles: activation domains balance hydrophobic motifs and acidic residues, acidic residues better promote exposure when adjacent to the motifs; within the motifs, the choice between aromatic and leucine residues is constrained by the structure of the coactivator interaction surface. Taken together, our results support the Acidic Exposure Model and provide a framework for unifying the roles of acidity, hydrophobicity, and intrinsic disorder in acidic ADs.

## Results

We investigated three key features of acidic ADs: acidic residues, hydrophobic motifs and disorder-to-order transitions. We designed sequence variants that systematically added and subtracted acidic residues or aromatic residues in seven ADs: VP16 (H1 region, 415-453), Hif1ɑ (AD2, 781-896), CITED2 (220-258), Stat3 (719-764), p65 (AD2, 521-551), p53 AD1 (1-40) and p53 AD2 (40-60) (Figure 1B). For each disordered region that folds into an alpha helix upon coactivator binding, we introduced proline or glycine residues, which suppress helicity (Figure 1–figure supplement 1). We measured 525 and 2991 variants in two experiments.

To test these designed variants, we developed a high-throughput method to assay AD variants in parallel in human cell culture (Figure 1C, Figure 1–figure supplement 2). We engineered a cell culture system with a synthetic TF that binds and activates a genome-integrated GFP reporter. Each cell receives one AD variant marked by a unique DNA barcode integrated into the same genomic “landing pad,” reducing the effects of genomic position on expression (Maricque et al., 2018). AD variants that drive different levels of GFP expression are separated by Fluorescent Activated Cell Sorting (FACS), and the barcodes in each sorted pool are counted by deep sequencing (Kinney et al., 2010; Sharon et al., 2012; Staller et al., 2018). The assay is reproducible, quantitative and resolves 9-10 levels of activity (Figure 2, Figure 2–figure supplement 1).

**Figure 2.**
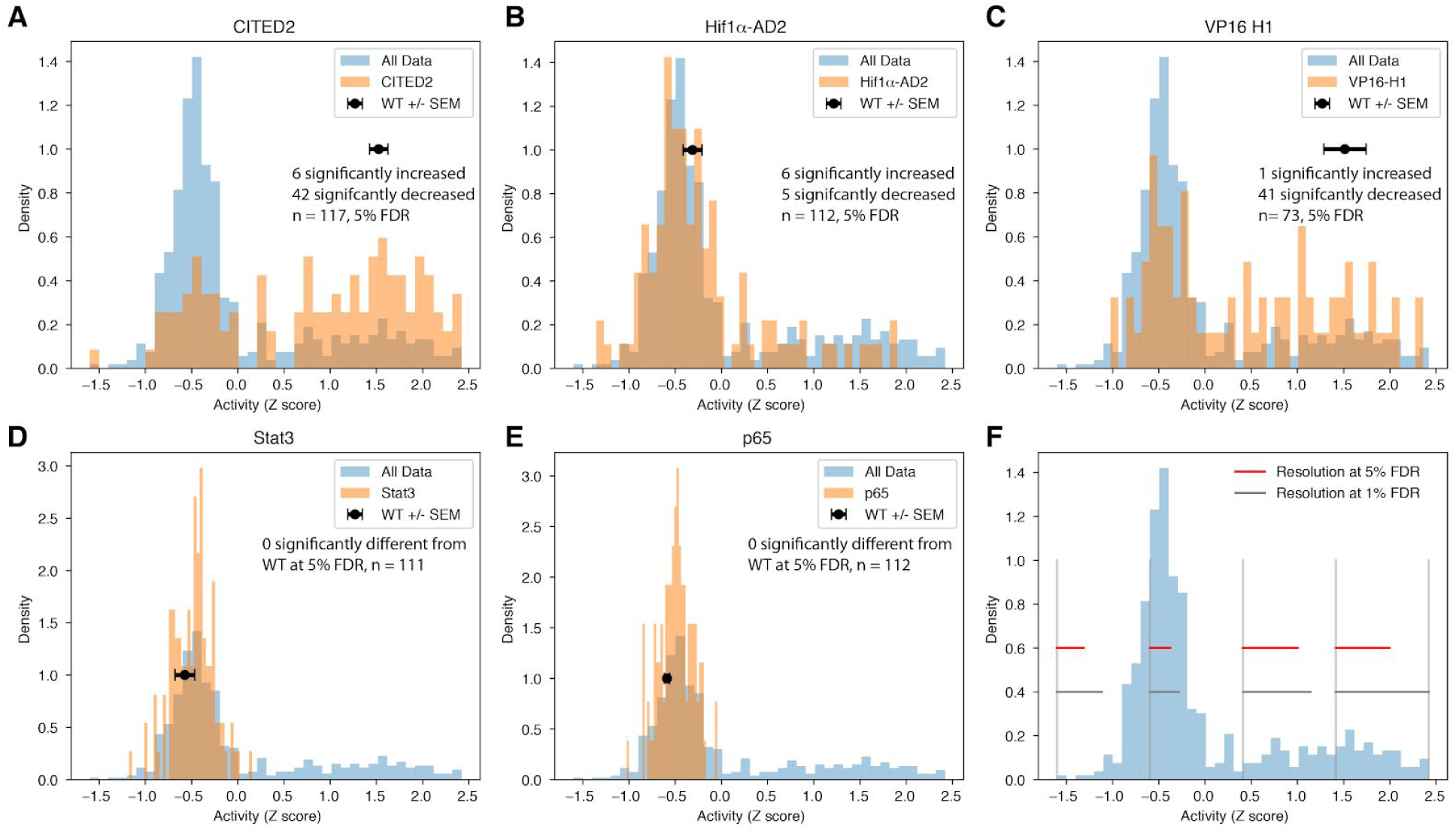
Rationally designed variants increase and decrease activation domain activity. Histograms of the activities of all variants (blue) and sets of specific AD variants (orange). The mean and SEM of each WT AD is shown in black. A) CITED2 and C) VP16 begin with high activity and designed variants increased or decreased activity. B) Hif1ɑ started with modest activity and designed variants increased and decreased activity. D-E) In Stat3 and p65, none of the designed variants significantly changed activity. F) The resolution of the method at 5% FDR (red) and 1% FDR (dark gray) in each quartile is shown (see methods). For 5% FDR, if we average the four red bars, we can resolve 9.3 levels of activity. If we apply each red bar to its respective quartile, we can resolve 10.4 levels of activity.

We tested four predictions of our Acidic Exposure Model. 1) Removing hydrophobic motifs will reduce AD activity. 2) Removing acidic residues (net negative charge) will reduce AD activity. 3) Adding acidic residues will increase AD activity. 4) Adding aromatic residues will increase AD activity.

### Prediction 1: Removing hydrophobic motifs will reduce AD activity

We confirmed that hydrophobic motifs make large contributions to AD activity (Cress and Triezenberg, 1991; Jackson et al., 1996; Lin et al., 1994). Removing aromatic and leucine-rich motifs decreased activity of all ADs (Figure 3A, 3B, Figure 3–figure supplement 1). Both known motifs (LPEL in CITED2, LPQL and LLxxL in Hif1ɑ, and LxxFxL in VP16 (Berlow et al., 2017; Regier et al., 1993)) and predicted motifs (Figure 1–figure supplement 1) contributed to activity. Aromatic residues made large contributions to activity in VP16 and both p53 ADs, as expected (Cress and Triezenberg, 1991; Lin et al., 1994), but made smaller contributions to activity in CITED2 and Hif1ɑ (Figure 4, Figure 4–figure supplement 1, Figure 4–figure supplement 2). These results confirm that AD activity requires hydrophobic motifs.

**Figure 3.**
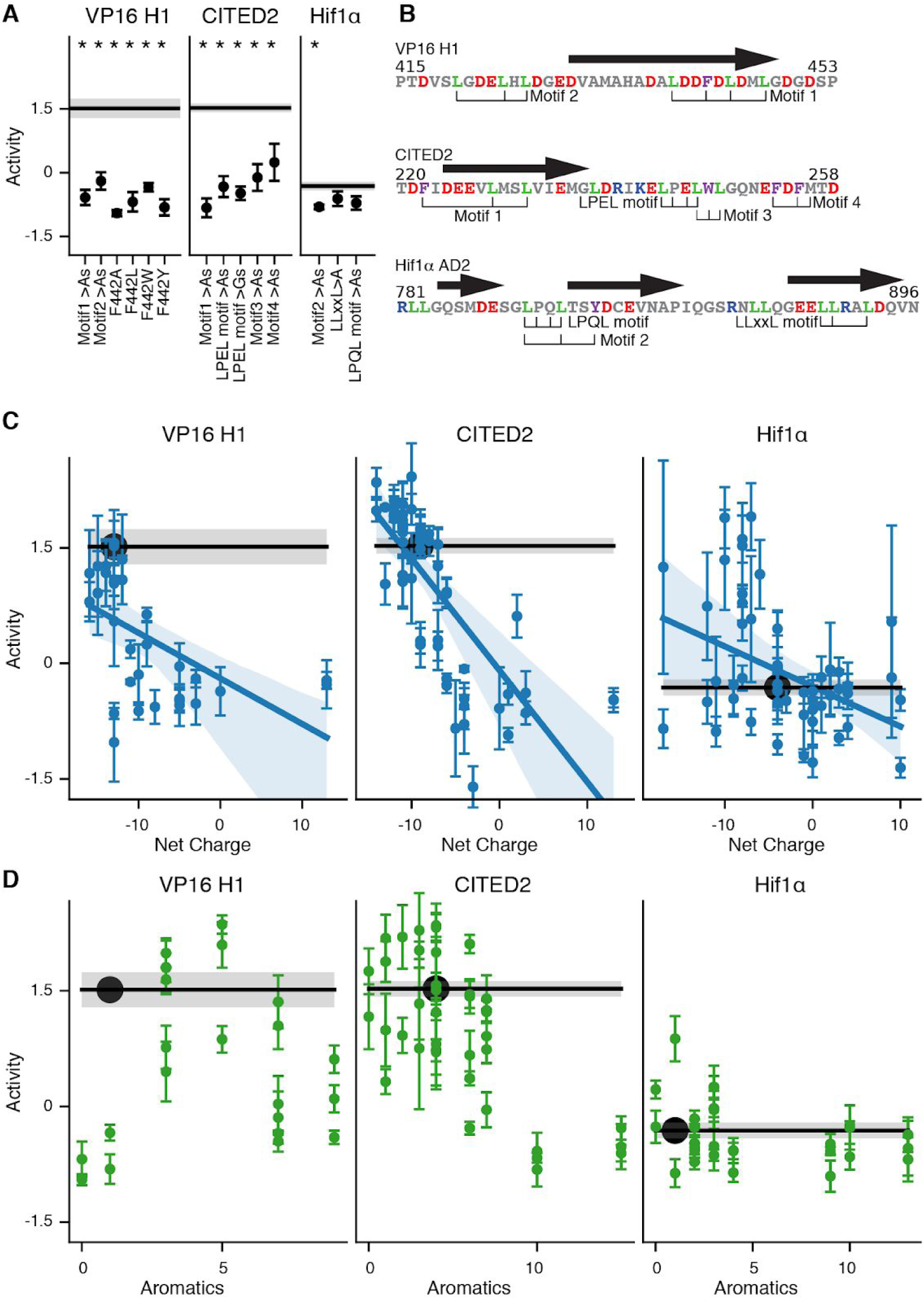
Acidic activation domains require hydrophobic motifs and acidic residues. A) Mutating motifs to alanine decreased activity. Mean and SEM shown. *, p<0.05. B) Motif locations and alpha helices (arrows). C) For variants designed to perturb net charge, the mean and SEM are plotted along with a linear regression and confidence interval. D) Variants that added aromatic residues increased or decreased activity. WT mean, black line and dot. SEM, gray box.

**Figure 4:**
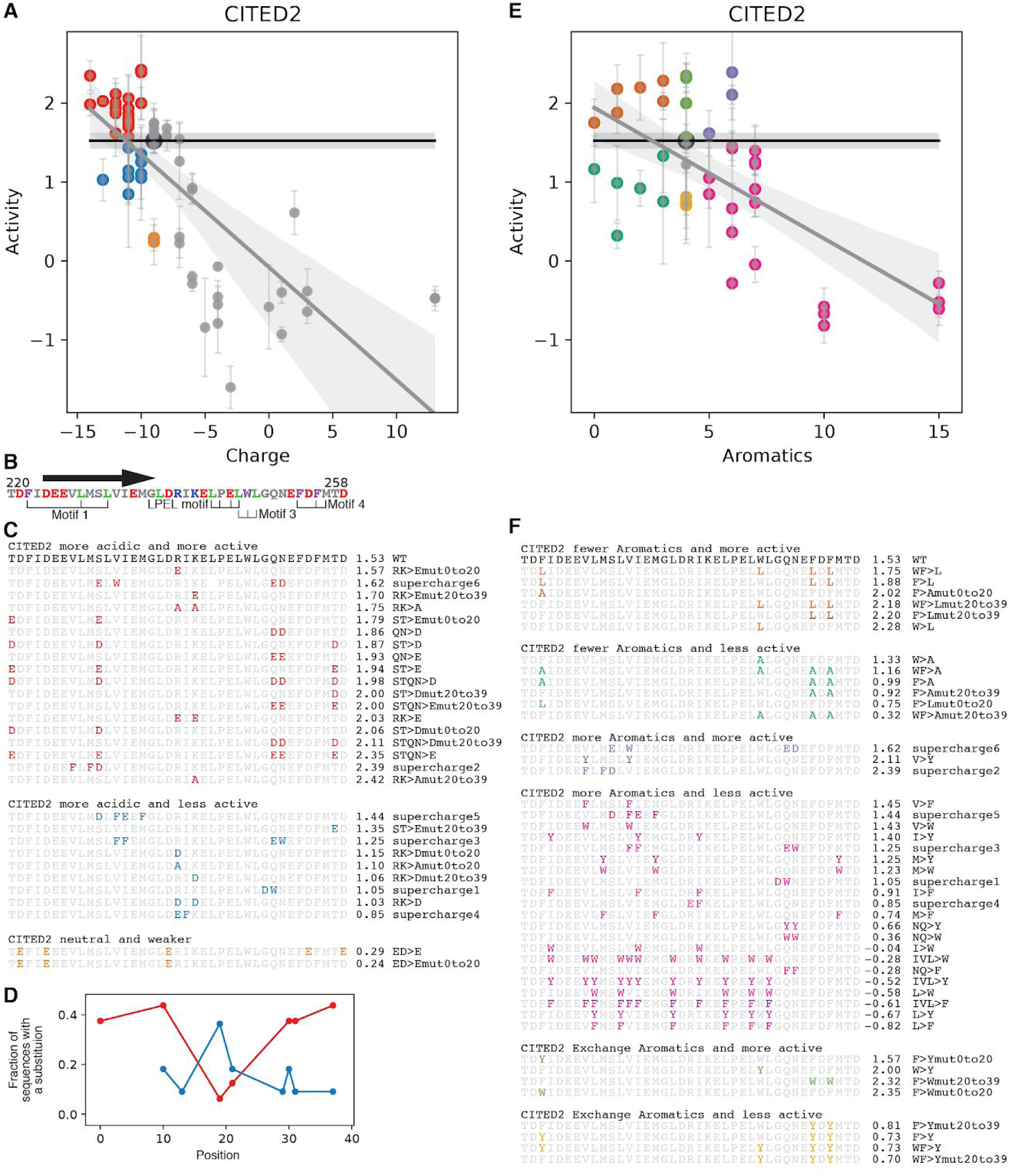
In CITED2 introducing acidic residues near hydrophobic motifs tends to increase activity. A) Variants that reduced net change and had increased activity are red. Variants that had reduced net charge and had WT or lower activity are blue. B) The location of the motifs and alpha helix (arrow). C) The sequences of variants colored in A. Red variants tend to add negative charge in the flanks, near the motifs. Blue variants tend to remove positive charges in the center or be ‘supercharge’ variants that add acidic and aromatic residues. Most electrostatically neutral substitutions had a small effect on activity, but two variants (orange) that substituted E for D in the N terminal region had reduced activity (discussed below). D) For the red and blue sets of variants, the fraction of variants with a substitution at each position is shown. Adding negative charges in the flanks, near the motifs, increases activity while adding negative charge in the center did not. E-F) CITED2 does not require aromatic residues for activity. Variants with fewer aromatic residues and higher activity (brown) generally replaced aromatics with leucines. Variants with fewer aromatic residues and lower activity (green). In contrast to VP16, replacing all aromatics with alanine (WF>A) only mildly decreased activity. In nearly all cases, adding aromatic residues decreased activity (pink). Two of the three variants that added aromatic residues and increased activity were ‘supercharge’ variants that also added acidic residues. Substituting F’s with W’s can increase activity (lime green). Substituting the F’s with Y’s generally decreased activity (orange). In both panels, WT mean and SEM are shown with a black line and gray horizontal box. A linear regression line and CI are plotted.

### Prediction 2: Removing acidic residues (net negative charge) will reduce AD activity

Acidic residues were necessary for the activity of all ADs. For VP16, CITED2, p53 AD1 and p53 AD2, removing acidic residues (moving towards positive net charge) decreased activity (Figure 3C, Figure 3–figure supplement 1, Figure 3–figure supplement 2). For Hif1ɑ, four of five variants with significantly reduced activity added basic residues (Figure 3–figure supplement 3). Activity was better explained by net charge than by the identities of the added acidic and basic residues: 1) removing negative residues or adding positive residues had similar effects, 2) adding lysine or arginine (both positively charged) had similar effects (Figure 3–figure supplement 4). All tested ADs required a net negative charge.

### Prediction 3: Adding acidic residues will increase AD activity

Adding negatively charged residues frequently increased the activity of CITED2, Hif1ɑ and p53 AD1 (Figure 3C, Figure 3–figure supplement 1). In Hif1ɑ, all variants with significantly increased activity decreased net charge (Figure 3–figure supplement 3). The location of added acidic residues determined whether activity increased: in CITED2 and Hif1ɑ, variants that added acidic residues adjacent to an aromatic or leucine residues were more likely to have increased activity (Figure 4, Figure 4–figure supplement 1, Figure 4–figure supplement 2). For the most acidic ADs, VP16 and p53 AD2, adding acidic residues rarely increased activity (Figure 3C, Figure 3–figure supplement 1, Figure 4–figure supplement 2). Once again, activity was better explained by net charge than the identity of the added residues: 1) adding aspartic acid or glutamic acid (both negatively charged) had similar effects and 2) removing basic residues was similar to adding acidic residues (Figure 3–figure supplement 4). Adding acidic residues caused two responses: increased activity or near WT activity.

### Prediction 4: Adding aromatic residues will increase AD activity

Adding aromatic residues can increase or decrease activity (Figure 3D). For VP16, adding 1-4 aromatic residues increased activity in the majority of variants, but adding more aromatic residues always decreased activity. For both p53 ADs, adding aromatics frequently increased activity (Figure 3–figure supplement 1). In contrast, adding aromatic residues to CITED2 and Hif1ɑ nearly always decreased activity (Figure 3D). Adding aromatic residues caused two opposite responses: increased activity or decreased activity. These two opposite responses did not match our prediction, but instead revealed a regime of the model we had not detected in yeast.

The Acidic Exposure Model can explain why the two responses to adding acidic residues mirror the two opposite responses to adding aromatic residues. In the model, adding acidic residues will increase AD activity only when there are hydrophobic motifs that can be further exposed. Once the hydrophobic motifs are maximally exposed adding more acidic residues will not increase activity. In contrast, adding more aromatic residues can increase activity only when there is excess exposure capacity, and adding too many aromatic residues eventually reduces activity, because they overwhelm the acidic residues and disorder to drive collapse. CITED2 has the most aromatic residues and its activity can be increased by adding acidic residues but not by adding aromatic residues. CITED2 has excess hydrophobic residues and activity is limited by exposure capacity (negatively charged residues). VP16 is the most acidic AD and its activity can be increased by adding aromatic residues but not by adding acidic residues. VP16 activity first increases then decreases as the number of added aromatic residues increases. Under the Acidic Exposure Model, VP16 has excess exposure capacity and thus, its activity increases as aromatic residues are added until the aromatics overwhelm the exposure capacity and drive chain collapse. This increase and decrease in activity has also been seen in synthetic ADs in yeast (Sanborn et al., 2020).

We further explored the Acidic Exposure Model by running all-atom Monte Carlo simulations of all VP16 and CITED2 variants (Methods) (Staller et al., 2018; Vitalis and Pappu, 2009). These simulations are optimized for disordered proteins and can detect the diversity of conformational ensembles. We found support for the idea that adding aromatics can overwhelm the exposure capacity of acidic and disorder promoting residues: introducing aromatic residues leads to more collapsed ensembles in the simulations (Figure 4–figure supplement 3). Conversely, adding acidic residues leads to more expanded ensembles. In the Acidic Exposure Model, maximal activity requires a balance between the numbers of acidic residues and hydrophobic motifs.

### A larger role for leucine residues in human ADs

The Acidic Exposure Model extends from yeast to human cells with one elaboration: a larger role for leucine residues in human cells. In yeast, we focused on the central AD of Gcn4, where aromatic residues made large contributions to activity while leucine and methionine made smaller contributions (Staller et al., 2018). Other groups have also observed that aromatic residues contribute more to AD activity than smaller hydrophobic residues (Erijman et al., 2020; Jackson et al., 1996; Ravarani et al., 2018; Sanborn et al., 2020). In human cells, we found three pieces of evidence that leucine residues make large contributions to AD activity. First, some motifs contained only leucine residues (Figure 3B). Second, replacing aromatic residues with leucine residues increased activity in Hif1ɑ and CITED2 (Figure 4, Figure 4–figure supplement 1). Third, replacing leucine residues with aromatic residues frequently decreased activity (Figure 4, Figure 4–figure supplement 1). The increased role of leucine residues likely reflects interactions with the expanded set of coactivators present in human cells.

### Strong ADs balance hydrophobic and acidic residues

We found that strong AD activity requires a combination of aromatic and leucine (W,F,Y&L) residues and acidic residues. Plotting the number of W,F,Y&L residues against net charge separates high and low activity variants (Figure 5A). Although neither the W,F,Y&L count nor net negative charge is sufficient for activity, both are necessary. Counting only aromatic residues does not separate high and low activity variants as well as counting W,F,Y&L residues (Figure 5–figure supplement 1). This separation is also visible when we normalize by AD length or substitute W,F,Y&L count with hydrophobicity (Figure 5–figure supplement 1). There are points on this grid occupied by both strong and weak variants (Figure 5–figure supplement 1), indicating that composition is not the sole determinant of activity and that the arrangement of residues also matters. When we used machine learning classifiers to separate active and inactive variants, leucines made larger contributions to model performance than any individual aromatic residue (Figure 5–figure supplement 1 and Figure 5–figure supplement 2). Our results suggest that the balance between W,F,Y&L and acidic residues is critical for AD activity.

**Figure 5.**
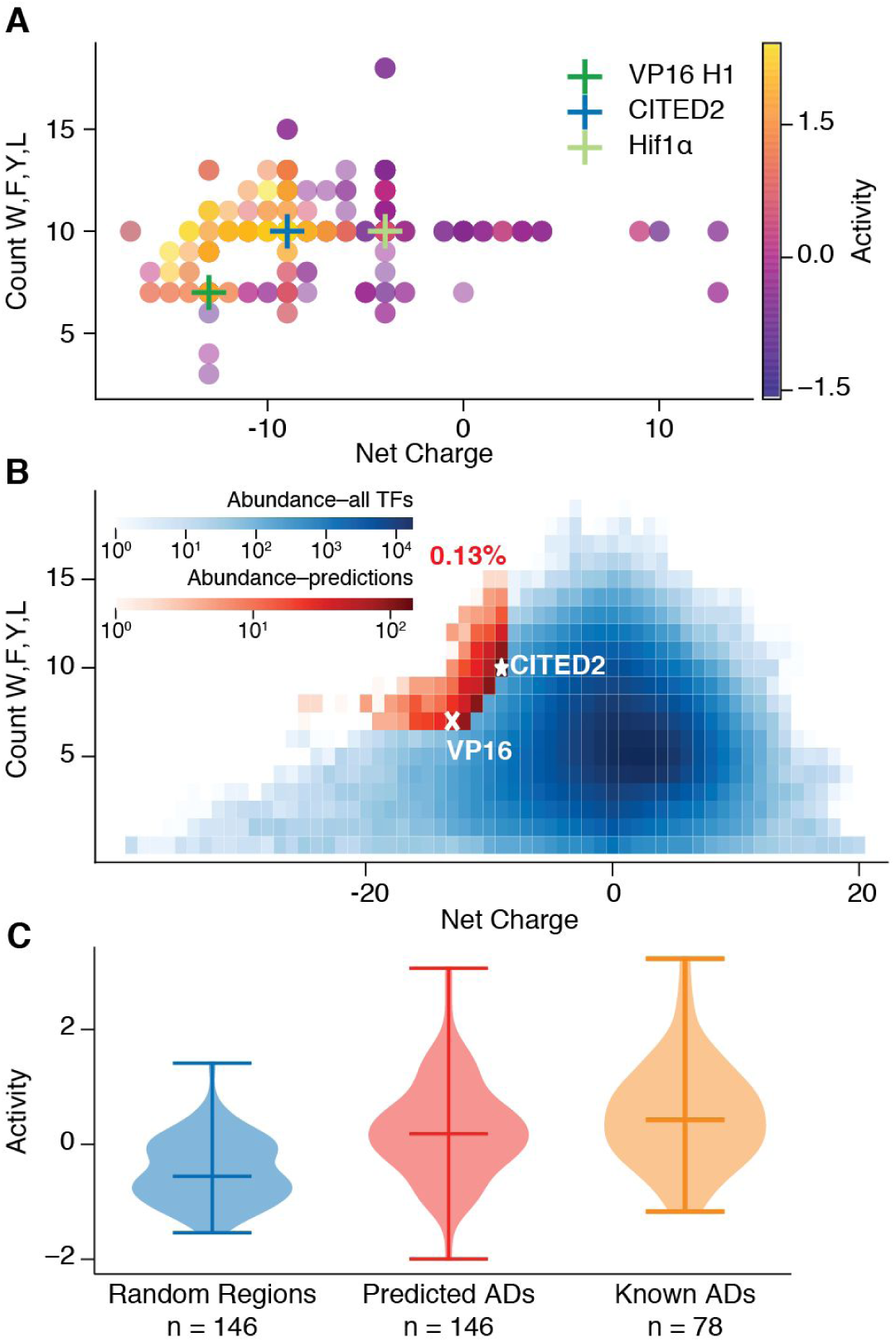
Strong activation domains are acidic and contain many W,F,Y&L residues. **A)** The location of each point indicates the properties, while color indicates activity. All variants of VP16, CITED2 and Hif1ɑ are plotted. **B)** A heatmap of all 39AA tiles from human TFs. The pixel location indicates the net charge and W,F,Y&L count, while the blue intensity indicates the number of tiles with that combination. Only 0.13% of tiles (red, rescaled heatmap) are as extreme or more extreme than VP16 (x) and CITED2 (*). **C)** TF regions spanned by the red tiles (red, n = 146) are more likely to have AD activity than random regions (blue, n=146). Most, but not all, published ADs (orange, n = 78) have high activity in this assay.

### Predicting ADs on human transcription factors

As a further test of the Acidic Exposure Model, we examined whether the balance of acidic and W,F,Y&L residues predicts known and new ADs in human TFs. For a third of human TFs, the only annotated domain is the DNA binding domain (Lambert et al., 2018) and only 8% of TFs have an AD annotated in Uniprot (Methods). *In silico*, we broke the protein sequences of 1608 TFs (Lambert et al., 2018) into 39 residue tiling windows (“tiles”), and for each tile we calculated the net charge and counted W,F,Y&L residues. Tiles that have both the net charge and hydrophobicity of strong ADs are rare: only 0.02% and 0.03% of tiles were as extreme or more extreme than VP16 or CITED2, respectively. Interpolating between these ADs yields 0.13% of tiles (n = 1139, Figure 5B, red), which combine to predict 144 ADs distributed across 136 TFs. These predicted ADs overlap with 17 Uniprot ADs - far more than expected by chance (p<1e-5 in permutation tests). In addition, 11 predicted regions overlap 10 published ADs that are not in Uniprot (p<1e-5 in permutation tests), including the N terminal AD of c-Myc (Andresen et al., 2012) and the Zn473 KRAB domain which has been recently shown to be an AD (Tycko et al., 2020). The high degree of overlap between our predicted ADs and literature-validated ADs motivated us to test the remaining predictions experimentally.

### Testing predicted ADs

We tested the predicted ADs and found that our composition model accurately predicted ADs in the human proteome (Figure 5C). In a new library, we tested 150 predicted regions (we split long regions to meet synthesis limits), 150 length-matched random regions and 92 published ADs (methods). We recovered 146 predicted ADs, 146 random regions and 78 published ADs. With the random regions, we built an empirical null distribution for the prevalence of ADs on human TFs, choosing the 95th percentile as the threshold for strong AD activity. At this high threshold, 39.7% (58/146) of our predicted regions are strong ADs and 51.3% (40/78) of published ADs are highly active. As a threshold for weak AD activity, we chose the median of the random regions, and found that 85.6% (125/146) of predicted ADs are active and 92.3% (72/78) of published ADs are active. This analysis demonstrates that our predictor is sufficient to identify known ADs and accurately predict new ones.

### Structural constraints of AD-coactivator interfaces

The AD predictor required counting leucine residues, prompting us to investigate the molecular role of leucines in human ADs. Summarizing the activities of all variants with a substitution at each position highlighted how mutating leucines decreased activity (Figure 6A, S12). Moreover, the residues with the lowest mean activities when mutated all point towards the coactivator surface in NMR structures of CITED2 and Hif1ɑ (Figure 6B, Figure 6–figure supplement 1). These same positions are also the most conserved positions of these ADs (Figure 6–figure supplement 2). These results support the hypothesis that the structural constraints of the interaction interfaces impose negative selection on the residues that make contact.

**Figure 6.**
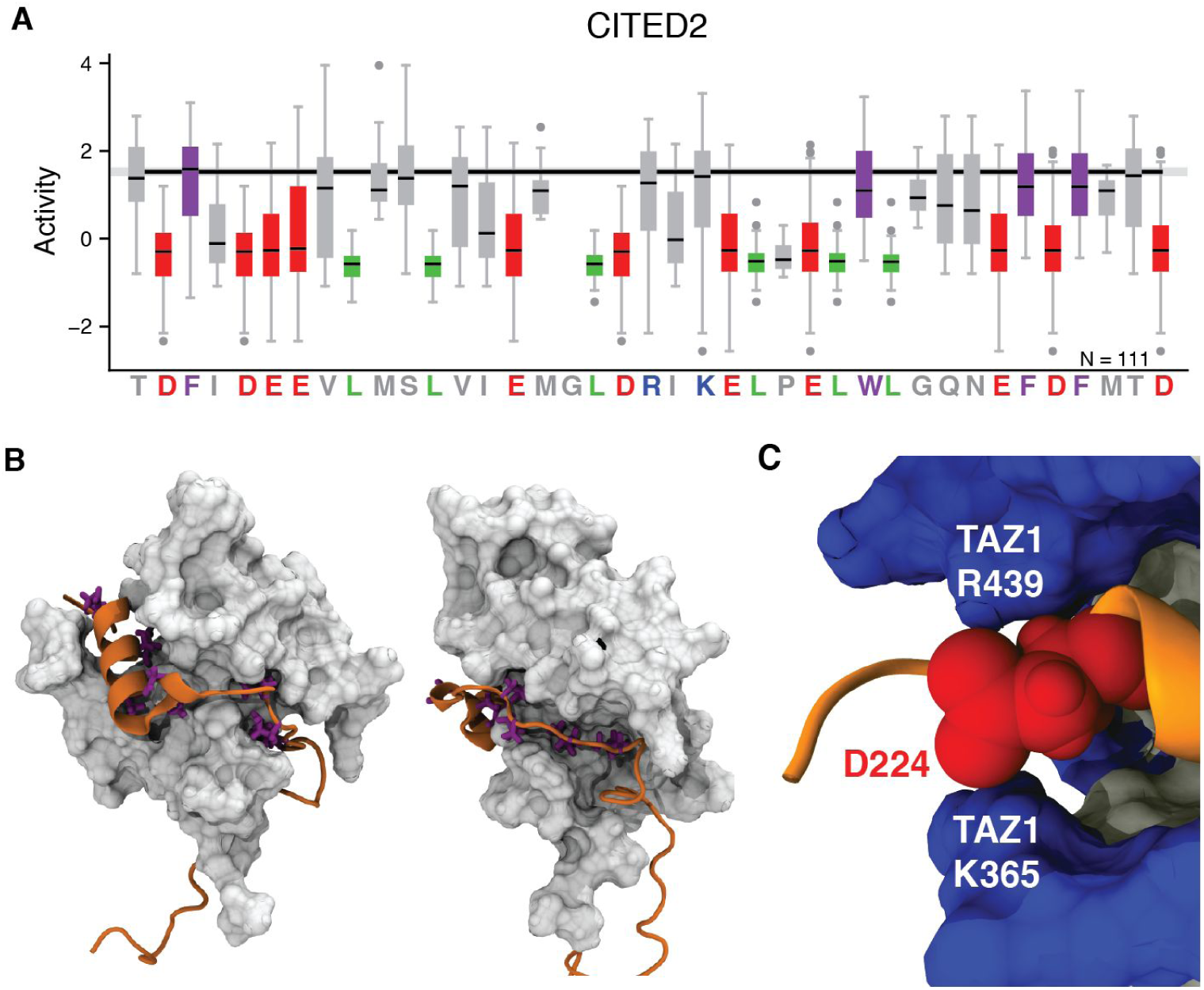
Rational mutagenesis detects the structural constraints of AD-coactivator interactions. A) The distribution of activities for all variants that change each position of CITED2. Acidic (red), aromatic (purple) and leucine (green) residues. Medians, gray bars. Outliers, gray dots. B) The residues of CITED2 (orange) with lowest mean positions in A (pink) point towards the TAZ1 coactivator (white, 1R8U). C) D224 (red) of CITED2 is sandwiched between the narrowest point of the basic rim (blue) of the binding canyon of TAZ1. See Figure 6–figure supplement 2 for snapshots of all 20 structures in 1R8U.

The mechanism by which leucine residues make large contributions to activity is exemplified by the CITED2 interaction with the TAZ1 domain of CBP/p300. TAZ1 has a canyon with a hydrophobic floor and basic rim that tightly embraces the compact alpha helix of CITED2 (Figure 6B, Figure 7B). The leucine residues on CITED2 interact with the hydrophobic canyon floor and the acidic residues interact with the basic canyon rim. This tight structural constraint explains the activities of many variants. Replacing leucines with aromatics decreases activity because the larger side chains do not fit in the canyon. Disrupting the helix folding, either by adding two prolines or shuffling the sequence (See simulations, Figure 6–figure supplement 3), causes expansion, reducing activity. Conversely, the 2xGlycine variant increased activity because it had smaller side chains and retained helicity. This structural constraint also explains why a conservative substitution, replacing aspartic acid residues (D) with larger glutamic acid residues (E), reduced activity: the D224 side chain sits between the narrowest point of the basic canyon rim, sandwiched between R439 and K365 of TAZ1, and the D244E substitution impairs this fit (Figure 4A, Figure 6C, Figure 6–figure supplement 4). If the signature we see in CITED2 carries over to other ADs that bind this TAZ1 canyon, then our results, along with other studies (Diss and Lehner, 2018; Rollins et al., 2019; Schmiedel and Lehner, 2019), could predict this family AD-coactivator interactions.

**Figure 7:**
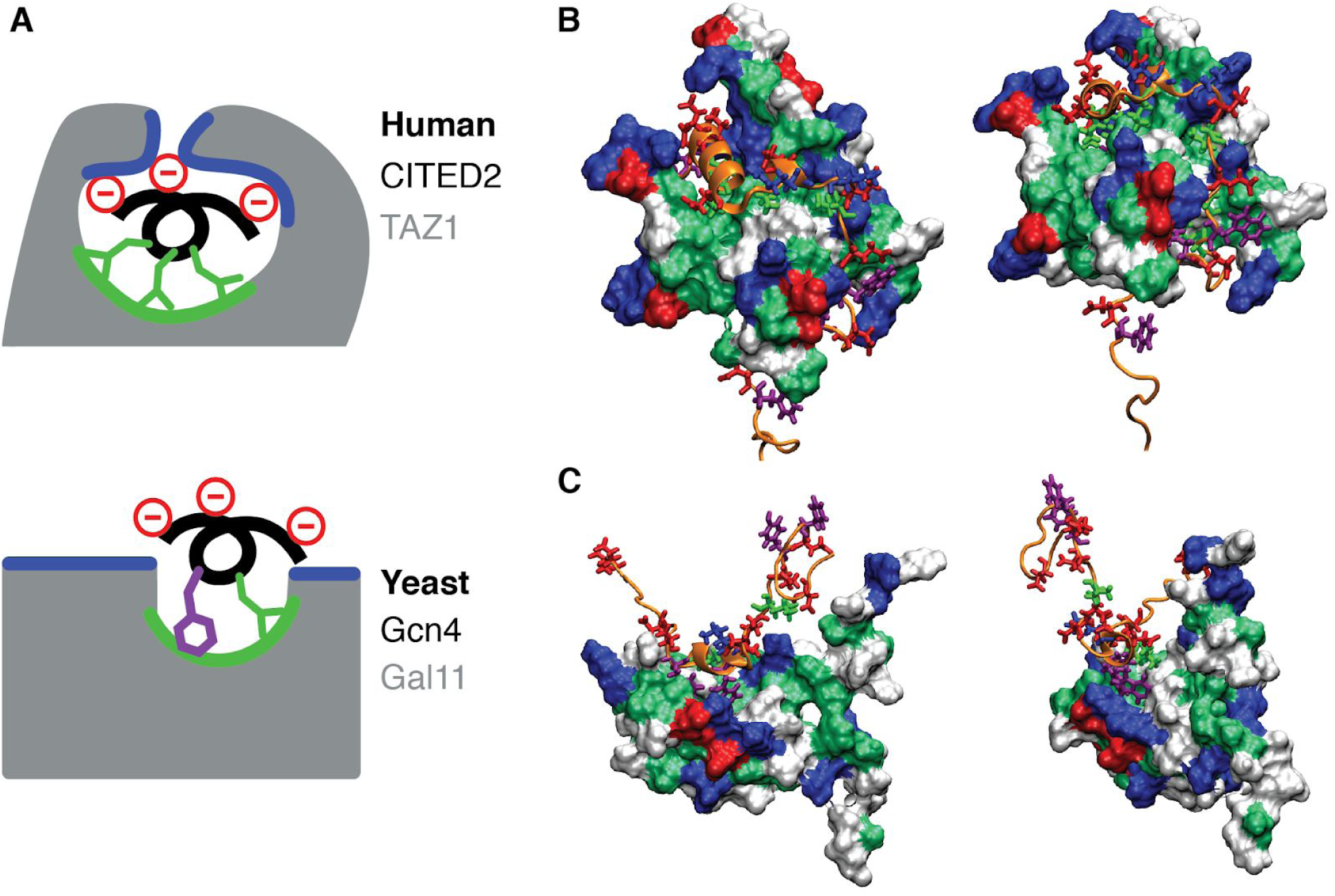
The structure of the coactivator AD-binding canyon constrains AD sequence. A) The CITED2 AD is inside the Taz1 canyon, a structural constraint that favors leucine residues. The yeast Gcn4 AD is outside the Med15/Gal11 canyon, enabling a fuzzy interaction that favors aromatic residues. B-C) Structures of the CITED2-TAZ1 and Gcn4-Med15 interactions. Both ADs fold into amphipathic alpha helices (orange) that present the hydrophobic residues (leucine, lime green or aromatic, purple) on one side and the acidic residues (red) on the other side. TAZ1 and Med15 each contain an AD binding canyon with a hydrophobic floor (dark green) and basic rim (blue). B) The large TAZ1 canyon embraces the compact CITED2 helix. Structure 1 of 1R8U. C) On Med15, the canyon is shallow, the Gcn4 alpha helix remains outside and inserts side chains. Structure 10 of 2LPB.

Contrasting the CITED2-TAZ1 interaction with the Gcn4-Med15 interaction explains why aromatic residues make large contributions to activity in Gcn4 (Berlow et al., 2017; Brzovic et al., 2011). Both ADs fold into alpha helices and both coactivators contain an AD binding canyon with a hydrophobic floor and basic rim (Figure 7). On TAZ1, the canyon is large and the CITED2 alpha helix is fully engulfed. Leucines fit this structure better than aromatics because they are smaller and promote helix formation (Pace and Scholtz, 1998). On Med15, the canyon is shallow and Gcn4 only inserts side chains. Aromatics fit this structure better than leucines because they better reach the hydrophobic canyon floor. The Gcn4-Med15 interaction is fuzzy with few steric constraints, explaining why Gcn4 is poorly conserved and more tolerant of substitutions in mutagenesis experiments (Brzovic et al., 2011; Erijman et al., 2020; Staller et al., 2018; Warfield et al., 2014). The increased importance of leucine residues in human cells likely reflects the structural constraints imposed by an expanded repertoire of coactivators.

### Amino acid sequence grammar

Although composition is a key determinant of AD activity and sufficient for predicting ADs, we found four ways in which the arrangement of amino acids (i.e. sequence) is also important for function. First, for VP16 and CITED2, 80% of shuffle variants, which maintain composition but rearranged the order of amino acid residues in a region of the AD, disrupted activity (Figure 6–figure supplement 3, Figure 6–figure supplement 5). Second, for VP16 and CITED2, forming an alpha helix was necessary for activity because variants that disrupted helicity also reduced activity (Figure 6–figure supplement 3, Figure 6–figure supplement 5). Third, adding acidic residues adjacent to W,F,Y&L residues was more likely to increase activity (Figure 4, Figure 4–figure supplement 1, Figure 4–figure supplement 2). This result agrees with our work in yeast and two random peptide screens which found that [DE][WFY] dipeptides make large contributions to AD activity (Erijman et al., 2020; Ravarani et al., 2018; Staller et al., 2018). Interspersing acidic residues between W,F,Y&L residues more efficiently exposes the motifs.

## Discussion

In the Acidic Exposure Model, acidic residues and intrinsic disorder keep hydrophobic motifs exposed to solvent where they are available to bind coactivators. Acidic residues prevent local chain compaction through electrostatic repulsion and favorable free energies of solvation. This expansion exposes leucine and aromatic residues to the solvent. Intrinsic disorder further reduces the entropic cost of exposing motifs (i.e. organizing water around hydrophobic residues) because fluctuating between solvent exposed and solvent protected conformations can lower the average cost compared to constant exposure. Acidity and intrinsic disorder are self-reinforcing because acidic residues promote disorder (Oldfield and Dunker, 2014). The Acidic Exposure Model explains why activation domains are both negatively charged and intrinsically disordered: the acidic residues and intrinsic disorder combine to keep aromatic and leucine-rich motifs exposed and available to bind coactivators.

Promoting exposure of hydrophobic motifs is compatible with the other known roles of acidity and disorder in ADs. Acidic residues on ADs can interact with basic residues on coactivators and allow biphasic binding (Ferreira et al., 2005; Hermann et al., 2001). In specific cases, intramolecular electrostatic interactions between negatively charged ADs and positively charged DBDs can increase DNA binding specificity (Krois et al., 2018; Liu et al., 2008). We speculate that because acidic residues can repel the negatively charged DNA backbone, they may also reduce non-specific DNA binding. Intrinsic disorder gives ADs the flexibility to fold into different conformations when bound to different partners (Dyson and Wright, 2016). Intrinsic disorder and acidic residues together can increase the fraction of molecular collisions that lead to binding by enabling multiple folding trajectories (Kim et al., 2018; Kim and Chung, 2020). The Acidic Exposure Model is compatible with these biophysical mechanisms of AD-coactivator interaction.

A key property of ADs is the balance between the strength of their hydrophobic binding motifs and their capacity to keep those motifs exposed to solvent. Adding more hydrophobic residues to ADs will increase their activity so long as the intrinsic disorder and acidic residues can keep the excess hydrophobicity from collapsing the amino acid chain into an inactive conformation. ADs have high hydrophobicity and high acidity and their activity requires a balance between these two physical properties. We exploited this observation to create an AD predictor that scans for a high, but balanced composition of acidity and hydrophobicity. This predictor can be used to prioritize candidate acidic ADs on poorly characterized TFs in any metazoan genome.

Our results can be summarized with three design principles for mammalian ADs. First, strong ADs require a compositional balance between W,F,Y&L residues and acidic residues to keep hydrophobic binding motifs exposed to solvent. Second, acidic residues more effectively expose motifs when they are directly adjacent to the W,F,Y&L residues. Third, within hydrophobic motifs, aromatic residues are preferred when the binding pocket of the cognate coactivator is shallow while leucine residues predominate when coactivators have large binding canyons that engulf the AD. Going forward these rules will help refine computational models for predicting ADs, guide engineering of new ADs, predict AD-coactivator interactions, and inform models that predict the impact of genetic variation on AD function.

## Acknowledgments

We thank Minhee Park and Ahmed Khalil for sharing the synthetic DBD and promoter sequence ahead of publication; Kiersten Ruff, Avi Ramu and Nicole Rockweiler for bioinformatics help; Brittany Pioso for help with the cartoons; Jessica Hoisington-Lopez and MariaLynn Crosby for DNA sequencing. We thank members of the Cohen Lab for helpful discussions and comments on the manuscript.

## Author contributions

MVS and BAC designed the project and wrote the manuscript. MVS and ER collected data. MVS, ER and ASH analyzed data. RVP and BAC interpreted data. All authors edited the manuscript. The authors declare no competing interests.

### Funding

MVS was supported by Burroughs Wellcome Fund Postdoctoral Enrichment Program, American Cancer Society Postdoctoral Fellowship, and NIGMS K99131022. ER was supported on the McDonnell Genome Institute Opportunities in Genomics Research Program under grant NIH-R25HG006687. This work was supported by grants from the National Institutes of Health, NINDS 5R01NS056114 to RVP and NIGMS R01GM092910 to BAC, and the from the Children’s Discovery Institute CDI-LI-2018-765 to BAC.

## Data availability

All processed activity data is available in the supplementary materials. Simulation data is freely available upon request. The raw data will be available in the NIH GEO database. Our code will be available on Github at publication. The plasmids will be deposited in AddGene.

## Materials and methods

### Cell line construction

To engineer the K562 cell line we began with LP3 from (Maricque et al., 2018). First, we introduced a frameshift mutation to the GFP in the landing pad using Cas9 and a gRNA against GFP (AddGene 41819). Second, we integrated our reporter at the AAVS1 locus using Cas9, the SM58 SSBD2 T2 gRNA (from Shondra Miller, Washington University School of Medicine GEiC) and the pMVS184 reporter plasmid (Figure 6–figure supplement 6). Starting on Day 2, we selected with 1 ug/ml puromycin for three days. We tested candidate reporter clones with transfections of a synthetic TF (pMVS 223, Figure 6–figure supplement 7) carrying p53 AD1, choosing the clone with the largest dynamic range between baseline GFP and the brightest transfected cells.

Cells were grown in Iscove’s Modified Dulbecco’s Medium (IMDM) medium +10% FBS +1% Non Essential Amino Acids +1% PennStrep (Gibco). All transfections used the Invitrogen Neon electroporation machine using a 100 ul tip, 1.2 M cells and 5 ug of DNA.

### Rational Mutagenesis

The sequences of all 525 VP16, Hif1ɑ, CITED2, Stat3 and p65 variants are listed in Supplemental Dataset 1. “Hand Designed” variants are shown in Figure 1–figure supplement 1. The systematic mutagenesis added and removed charged residues or aromatic residues. Net charge of ADs was changed in two ways: subsets of charged residues were changed to each of the four charged residues and alanine, or subsets of polar residues were changed to charged residues. Aromatic residues were changed to alanine, leucine or other aromatic residues, and aromatic residues were added by replacing leucine, isoleucine, alanine, methionine, and valine residues.

The “Hand Designed” p53 AD variants contained the same systematic mutations and more hand designed variants listed in Supplemental Dataset 2. The p53 mutagenesis also included a deep mutational scan, a double alanine scan and some sequences from orthologous TFs. Activity values of each dataset are normalized separately.

### Plasmid Library Construction

The plasmid sequences for the GFP reporter (Figure 6–figure supplement 6) and synthetic TF (Figure 6–figure supplement 7) are in Supplemental Dataset 6. Both will be available from Addgene.

We designed the AD variants as protein sequences and reverse translated using optimal human codons. We attached each variant to 28 unique 12 bp FREE barcodes (Hawkins et al., 2018). WT ADs had 84 barcodes each. We added PCR primers at the start and end. Between the AD and the barcode are BamHI, SacI and NheI restriction sites. For ADs that were less than 46AA, we added random filler DNA between the BamHI and SacI sites. We ordered 14968 unique 217 bp ssDNA oligos from Agilent.

We cloned the AD variant library by HiFi assembly. We added plasmid homology to the ssDNA oligos by PCR, yielding a 232 bp product, with 4 cycles, Q5 polymerase, 0.5 pmol template and 8 reactions. We digested the pMVS223 backbone with AflII, XhoI and KpnI-HF and gel purified it. Each assembly had 100 ng of backbone and 5x molar ratio of insert. We electroporated bacteria and collected ∼20 million colonies. We checked the library with paired end Illumina sequencing. We recovered 98.7% of our barcodes and all AD variants. For the second step of library cloning, we digested the library and pMVS223 with BamHI-HF and NheI-HF, and inserted the synthetic TF by T4 ligation. We electroporated bacteria and collected 400K colonies. We recovered 93% of designed barcodes and all ADs.

The p53 library was constructed in the same way with 5 barcodes per variant, 30 for WT AD1 and 25 for WT AD2. We collected 4M colonies after step one and 26.2M after step two. We recovered 14355 of 14998 designed barcodes and 2990 of 2991 designed ADs. In this work we used data from both WT ADs and 171 hand designed variants.

All restriction enzymes, HiFi mix, and competent bacteria were purchased from NEB. Library Maxipreps were performed using the ZymoPURE II Plasmid Maxiprep Kit (Zymo).

### Plasmid Library Integration and measurement

In each transfection, we used 1.2 M cells, 2 ug of CMV-CRE (Maricque et al., 2018) and 3 ug of Plasmid Library. We transfected 102 M cells in 86 transfections split into 22 flasks. The next day, we began selection with 400 ng/ml G418 for 10 days. On Day 11 we performed magnetic enrichment of live cells (MACS). We combined flasks 1-5 into biological replicate 1, flasks 6-10 into biological replicate 2, flasks 11-15 into biological replicate 3, and flasks 16-22 into biological replicate 4. On Day 12, we added 1 uM ß-estradiol.

On Day 15 we sorted cells on a Sony HAPS 2 at the Siteman Cancer Center Flow Cytometry Core. We set an ON/OFF threshold for GFP as the 90th percentile of the uninduced population. The lowest bin was the bottom 50% of the OFF population. The ON region was split into 3 bins with equal populations. For each replicate, we collected 750K cells in each of the four bins. We noted the median fluorescence of each bin and used that number to calculate activity below. The dynamic range of the measurement is determined by the fluorescence values of the dimmest and brightest bin.

### Barcode amplicon sequencing libraries

Genomic DNA was collected using the Qiamp DNA Mini kit (Qiagen). We performed 8 PCRs on each sample. The sequencing libraries were prepared in 2 batches: Batch 1 contained biological replicates 1-3 and Batch 2 contained biological replicate 4. We did 25 cycles with NEB Q5 polymerase using CP36.P10 and LP_019 primers. We pooled the PCRs, cleaned up the DNA (NEB Monarch), quantified it, digested the entire sample with NheI and EcoRI-HF (NEB) for 90 minutes and then ligated sequencing adaptors with T4 ligase (NEB) for 30 min. We used 4 ng of this ligation for a 20 cycle enrichment PCR with Q5 and the EPCR_P1_short and EPCR_PE2_short primers. We sequenced each Batch on a NextSeq 500 1×75 High Output run. Each biological p53 replicate was sorted on a different day, so each sequencing library was sequenced separately with a NextSeq 500 1×75 High Output run.

CP36.P10 ctcccgattcgcagcgcatc

LP_019 GCAGCGTATCCACATAGCGTAAAAG

EPCR_P1_short AATGATACGGCGACCACCGAG

EPCR_PE2_short CAAGCAGAAGACGGCATACGAGAT

To assess the number of integrations in each experiment, we saved 1 ml of culture (0.5-1M cells) from each flask (4 transfections) before the magnetic enrichment for live cells (Day 11). We extracted gDNA, amplified barcodes, sequenced. We identified 96,000 unique integrations, an underestimate. In the sorted samples we recovered 14015 barcodes (93.6% of designed) total, 7164 in all four replicates and 10798 in three or more replicates. All ADs were present in all replicates.

### Data processing

We demultiplexed samples using a combination of Index1 reads and Read1 inline barcodes. Using perfect matches, we counted the abundance of each FREE barcode in each sample. We normalized the read counts first by the total reads in each sample and then renormalized each barcode across bins to create a probability mass function. We used the probability mass function and the median GFP fluorescence of each bin to calculate the activity of each barcode. To remove outlier barcodes, we found all barcodes for an AD, computed the activity of each barcode, computed the mean and variance of the set of barcodes and then removed any barcodes whose activity was more than two standard deviations away from the mean. We then took all the reads from all remaining barcodes, pooled them and recomputed activity. This approach led to one activity measurement for each biological replicate. We converted these activities into Z scores and computed the mean and standard error mean (SEM). The Z score normalization is nearly equivalent to mean centering the replicates (Pearson R^2^ =1). The Z score normalization was essential for combining the p53 data, because the fluorescence values differed between days. We used Z scores for both datasets for consistency. Pooling barcodes led to reproducible AD measurements (Figure 2–figure supplement 1B). Reproducibility improved with more barcodes (Figure 2–figure supplement 1C) and with more integrations (Figure 2–figure supplement 1D).

### Analysis

All analysis was performed in Jupyter Notebooks with python 2.7 and Matplotlib, seaborn, pandas, localcider, biopython, logomaker (Tareen and Kinney, 2020), scipy, statsmodels, sklearn, and ittertools. Colors from Colorbrewer (https://colorbrewer2.org/). AD sequence properties were calculated with localcider (Holehouse et al., 2017). To identify AD variants that were statistically significantly different from WT, we used a two sided t test and 5% FDR correction. We plotted the regressions with ‘regplot’ command in seaborn, using the ‘robust’ option for the confidence interval. To estimate the resolution of the assay (Figure 3D), for all pairs of ADs, we compared mean activities with a t-test, and corrected for multiple hypotheses 5% and 1% FDR. For each quartile, we averaged the ten smallest significant differences and plotted this mean difference Figure 1D and S3.

Structures were downloaded from the RSCB PDB (www.rcsb.org) and visualized with VMD (Humphrey et al., 1996). We normalized activity values to [0-1], mapped the values to the Beta column of the pdb file and visualized positions with normalized activity < 0.2 (Figure 4B, S16).

To summarize the effects of substitutions at each position (Figure 4, S16), we identified all variants that changed each position, collected the activity measurements from all biological replicates and created a boxplot. We excluded the shuffle variants.

To assess the conservation of each AD, we used the HMMER website (hmmer.org) Using the full protein sequences, we ran HMMER for 3 iterations, at which point, VP16 had converged with 47 sequences, CITED2 had converged with 446 sequences, and Hif1ɑ had not yet converged with 126K sequences. We took a screenshot of the logo of the HMM model for the alignment.

We performed ANOVA on composition, regressing activity of all four replicates against all 20 amino acids and a batch term. We iteratively removed parameters that were not significant. Separately, we trained a model with all dipeptides derived from [D,E,W,F,Y,L] and identified significant dipeptides. We added the significant dipeptides to the composition model and iteratively removed parameters that were not significant.

### All atom simulations

We ran all-atom, Monte Carlo simulations in the CAMPARI simulation engine (campari.sourceforge.net) using the ABSINTH implicit solvent paradigm (Vitalis and Pappu, 2009). This simulation framework is a well established approach to study the conformational ensembles of intrinsically disordered regions (Martin et al., 2016; Metskas and Rhoades, 2015; Vitalis and Pappu, 2009) and we have previously used it to study the Central Acidic AD of the yeast TF, Gcn4 (Staller et al., 2018). We simulated all VP16 and CITED2 variants. For Hif1ɑ, we simulated all hand designed variants and the WT sequence. All simulation data is available upon request.

For each variant, we ran ten simulations starting in a helix and ten starting in a random coil. In total we ran 4300 simulations. Each simulation had a pre-equilibration run of 2M steps. Then we began the real simulation with 10M steps of equilibration and the main simulation of 50M steps, extracting the conformation every 10K steps, yielding 5000 conformations per simulation. Simulation analysis was performed with the CAMPARItraj (ctraj.com) software suite. This software suite calculated helicity with the DSSP algorithm (Kabsch and Sander, 1983) and radius of gyration as the distribution of atoms in each confirmation without weighting by mass (Holehouse et al., 2017). The accessibility was calculated by rolling a 1.5 nm spherical marble around each confirmation and summing the solvent accessible surface area of the W,F,Y&L residues (Staller et al., 2018). To speed up this analysis accessibility was assessed every 20 confirmations.

### Machine Learning

The machine learning analysis was carried out in python with the sklearn package. We started with all variants of VP16, Hif1ɑ and CITED2 and then excluded the shuffle variants. The High Activity set (N = 122) had variants with a mean Z score above 0.5. The Low Activity set (N = 136) had variants with a mean Z score below 0. We normalized all parameters to be between [0,1]. We performed 5 fold cross validation and assessed model performance with the Area Under the Curve (AUC) of the Receiver Operator Characteristic (ROC).

### Predicting ADs in human TFs

We downloaded protein sequences from Uniprot for 1608 TFs (Lambert et al., 2018). For each TF, we created 39 AA tiling windows, spaced every 1 AA, yielding 881,344 tiles. For each tile, we computed the net charge (counting D,E,K&R) and counted W,F,Y&L residues.

We identified tiles that were as extreme or more extreme than VP16 and CITED2. We used a diagonal line to extrapolate between these ADs. The tiles predicted to cover ADs (Figure 4A, red pixels), fulfill 3 criteria:

(Charge < -9) AND (WFYL > 7) AND (((Charge+9)-(WFYL-10)) <= 0)

This algorithm identified 1139 tiles, 0.129% of the total. We aggregated overlapping tiles to predict 144 ADs on 136 TFs. To test these predictions, we used ADs annotated in Uniprot. We downloaded .gff files for the 1608 TFs from Uniprot. We used 4 regular expressions to search the “regions” column of the .gff files for “activation”, “TAD”, “Required for transcriptional activation” and “Required for transcriptional activation.” These searches yielded 110 unique ADs, including 7 proline rich ADs (>20% proline) and 3 glutamine rich ADs (>20% glutamine).

We used permutation tests to determine if our predictor was better than random. We randomly selected 136 TFs, randomly selected 144 length matched regions and determined how many overlapped the 110 known ADs. For the 4 TFs with 2 predicted ADs, we preserved the coupling between these lengths. In 100K permutations, we never observed more than 11 overlaps. 17 of our predicted ADs overlapped the 110 Uniprot ADs. We also applied this predictor to our AD variants in Figure 5–figure supplement 1.

### Testing Predicted ADs

We built a third plasmid library to test the predicted ADs. Due to DNA synthesis limits, we split long predicted regions and tested 150 regions of 39-76 residues. To create an empirical distribution for the prevalence of ADs on TFs, we included 150 length-matched regions randomly drawn from TF sequences(Lambert et al., 2018). We required that these random regions did not overlap our predicted ADs or Uniprot ADs. The 92 positive control ADs were drawn from: 36 hand-curated ADs (RegionType=Hand_Curated_ADs), 35 ADs from a published list (RegionType=Choi_2000_PMID_10821850)(Choi et al., 2000), 19 Uniprot domains that overlapped our predictions (RegionType=Uniprot), and 2 published synthetic DW or DF runs (RegionType=Controls)(Ravarani et al., 2018). We also included 3 KRAB domains from Uniprot, 22 mutant ADs and 26 regions tiling the human TF, Crx. Due to an error, we did not test the correct predicted region of AEBP1 (Q8IUX7) and tested 2 other regions instead. The full list of sequences and activities is included in Dataset 4. The ‘Known ADs’ in Figure 5C are flagged in the ‘Positive Controls’ column. The ‘Negative Controls’ column indicates mutant ADs.

The plasmid library was cloned in a similar manner as above. The oligos were ordered as a oPool from IDT. Oligos length varied. For each AD, we included one 9 bp ‘AD barcode’(Hawkins et al., 2018). During the second step of cloning, we added 6 Ns downstream of the synthetic TF by PCR, which became the ‘integration barcode.’ In principle, a different integration barcodes marks each plasmid integration event, analogous to a Unique Molecular Identifier in single cell RNA-seq protocols. The resulting ‘composite barcode’ contained a 6 bp ‘integration barcode’, the NheI restriction site and the 9 bp ‘AD barcode.’ 76 transfections were split into 3 biological replicates. G418 selection began on Day 1, magnetic separation was performed on Day 11, ß-estradiol induction began on Day 11 and cell sorting on Day 14. During the sequencing library preparation, we performed 24 PCRs for each gDNA sample.

After integrating the plasmid library into cells, we deeply sequenced the unsorted pool to build a ‘composite barcode’ table. This table contained 44077 composite barcodes with at least 10 reads in one of the 3 biological replicates. For all subsequent analysis, we matched reads to this table. We used perfect matches to designed ‘AD barcodes’ and combined reads for all ‘integration barcodes’ attached to each ‘AD barcode’ as described above: we first removed outliers and then combined read counts. The biological replicates contained 20850, 19758 and 21656 uniquely identifiable integrations. We designed 443 ADs, detected 434 in the plasmid library, and detected 431 integrated into cells. We required 5 or more unique integration barcodes in at least one replicate, yielding 407 ADs for downstream analysis.

## Supplemental Datasets

Dataset 1: All variants and activity measurements for the 5 AD library (VP16, CITED2, Hif1ɑ, Stat3 and p65).

Dataset 2: All variants and activity measurements for the p53 ADs.

Dataset 3: Predicted acidic ADs on human TFs.

Dataset 4: Sequences and activity measurements from testing predicted acidic Ads

Dataset 5: AD sequences and DNA barcodes for the 5 AD library.

Dataset 6: AD sequences and DNA barcodes for the p53 ADs.

Dataset 7: Plasmid sequences for pMVS184 and pMVS223.

---

**Figure 1–figure supplement 1:**
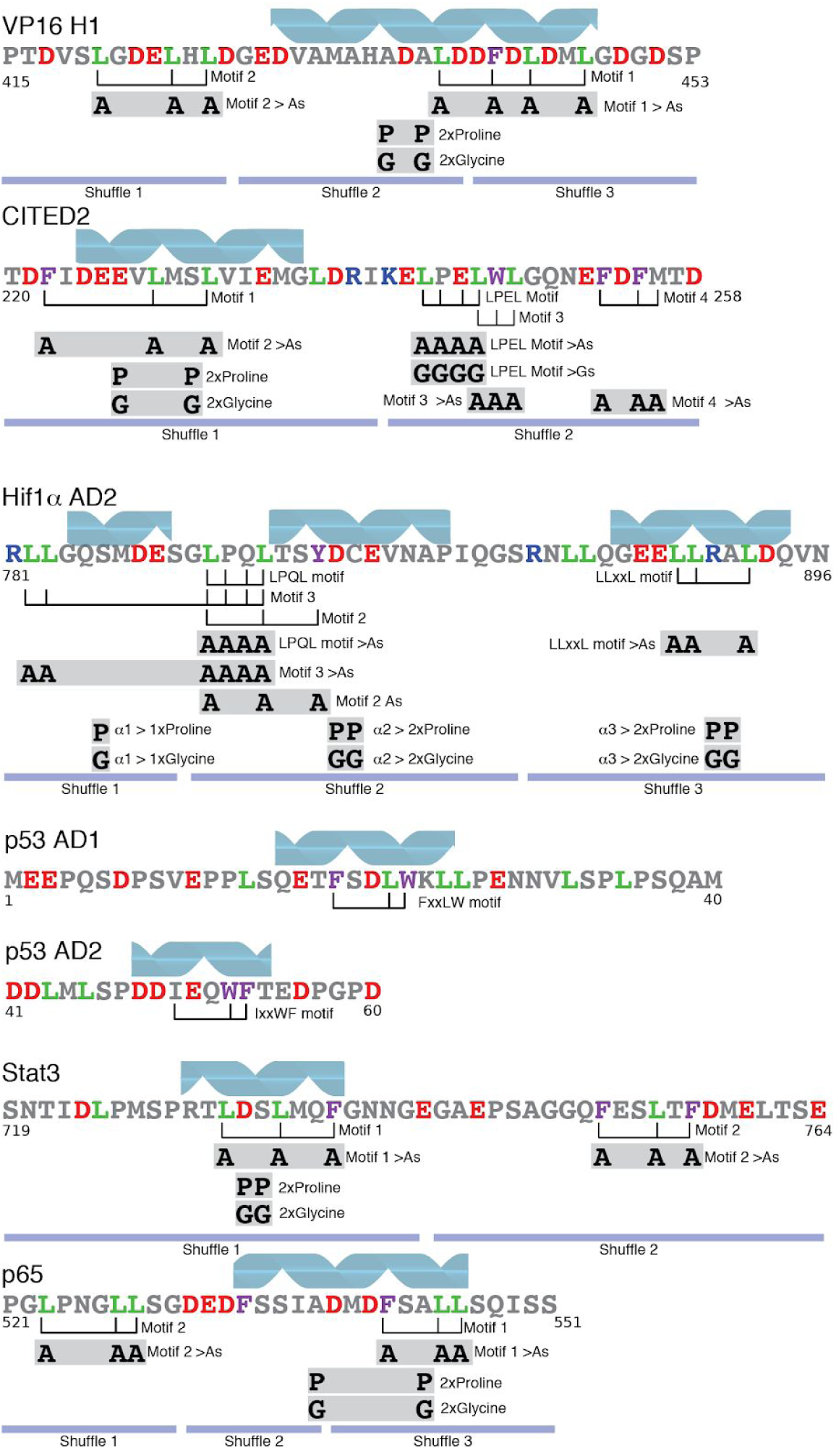
Cartoon of hand designed variants in each AD. Known and predicted alpha helices are indicated as blue helices (Berlow et al., 2017; Jonker et al., 2005; Krois et al., 2016; Lecoq et al., 2017; Wojciak et al., 2009). Hydrophobic motifs are indicated with brackets. Published motifs are LPEL in CITED2, LPQL and LLxxL in Hif1ɑ, LxxFxL in VP16, FxxLW in p53 AD1 and IxxWF in p53 AD2. (Berlow et al., 2017; Raj and Attardi, 2017; Regier et al., 1993). Variants that remove motifs are indicated as gray boxes. Variants designed to break the helices include the 2xProline and 2xGlycine substitutions. The regions that were randomly shuffled in each AD are indicated. The shuffle variants disrupt helices, disrupt motif grammar and test if the arrangement of residues contributes to activity.

**Figure 1–figure supplement 2:**
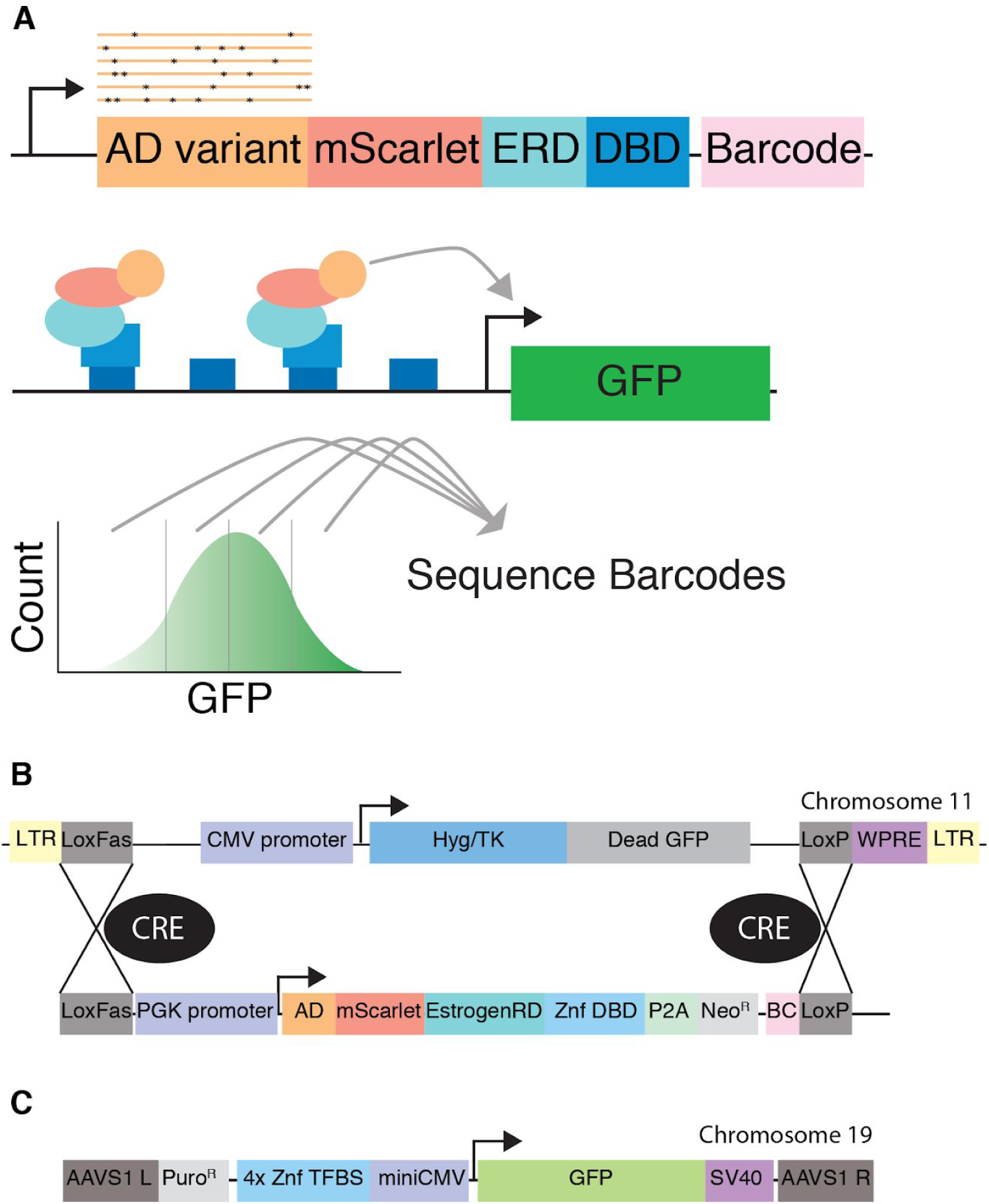
We developed a high throughput method for assaying AD activity in human cell culture. A) We clone a library of designed AD variants into a synthetic TF that contains an mScarlet red fluorescent protein, an estrogen response domain (ERD), a synthetic Zinc Finger DNA binding domain (DBD) and a barcode sequence in the 3’ UTR. When we induce nuclear localization with ß-estradiol, the synthetic TF enters the nucleus and activates a GFP reporter. The reporter has 4 binding sites for the DBD upstream of a miniCMV promoter. We use ‘Sort-seq’ to quantify activity of each variant (Kinney et al., 2010). We used landing pad #3 from (Maricque et al., 2018). B) Detailed cartoons of the Landing Pad and synthetic TF. Hyg/TK, hygromycin resistance and thymidine kinase fusion gene. WPRE, Woodchuck Hepatitis Virus Posttranscriptional Regulatory Element. LTR, lentivirus long terminal repeats. C) Cartoon of the reporter gene integrated at the AAVS1 locus. See Figure 6–figure supplement 6 and 7 for plasmid maps.

**Figure 2–figure supplement 1:**
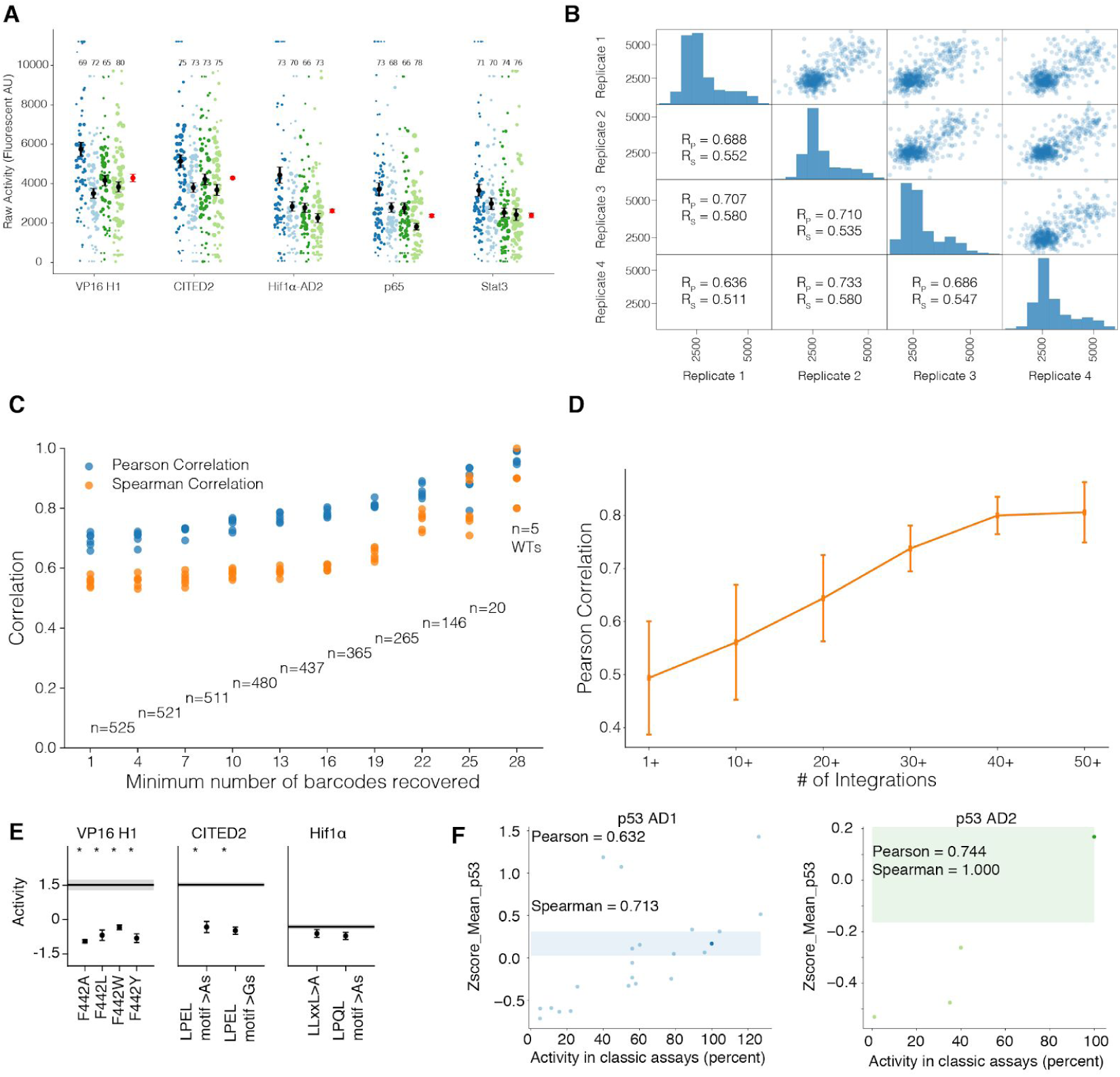
Validating the high throughput AD assay. A) Each WT AD was represented by 84 barcodes. The activity of each recovered barcode is plotted as a dot and the size of the dot is the square root of the number of reads for the barcode. For each AD, four biological replicates are shown and the number of barcodes recovered for each AD in each replicate is indicated. The mean and SEM calculated from combining data across all barcodes (red) agrees with the individual means (black). B) Reproducibility of AD measurements. Diagonal: histograms of AD activity measurements. Upper panels: reproducibility of ADs between biological replicate measurements. Lower panels: Pearson (R_P_) and Spearman (R_S_) correlation coefficients. C) The correlations between biological measurements improve as more barcodes are recovered. The 6 pairs of replicates are plotted. D) The correlations between biological replicates improve with the number of independent integrations. The mean and standard deviation of six pairs is shown. E) Variants of VP16, CITED2 and Hif1ɑ that have previously been shown to reduce activity reduce activity in our assay (Berlow et al., 2017; Cress and Triezenberg, 1991; Freedman et al., 2003; Regier et al., 1993). *, significant at 5% FDR. F) The activities of published variants of p53 are correlated with the activities we measure (Chang et al., 1995; Lin et al., 1994). WT p53 AD1, dark blue dot; SEM, light blue box. WT p53 AD2, dark green dot; SEM, green box (cutoff at the top).

**Figure 3–figure supplement 1:**
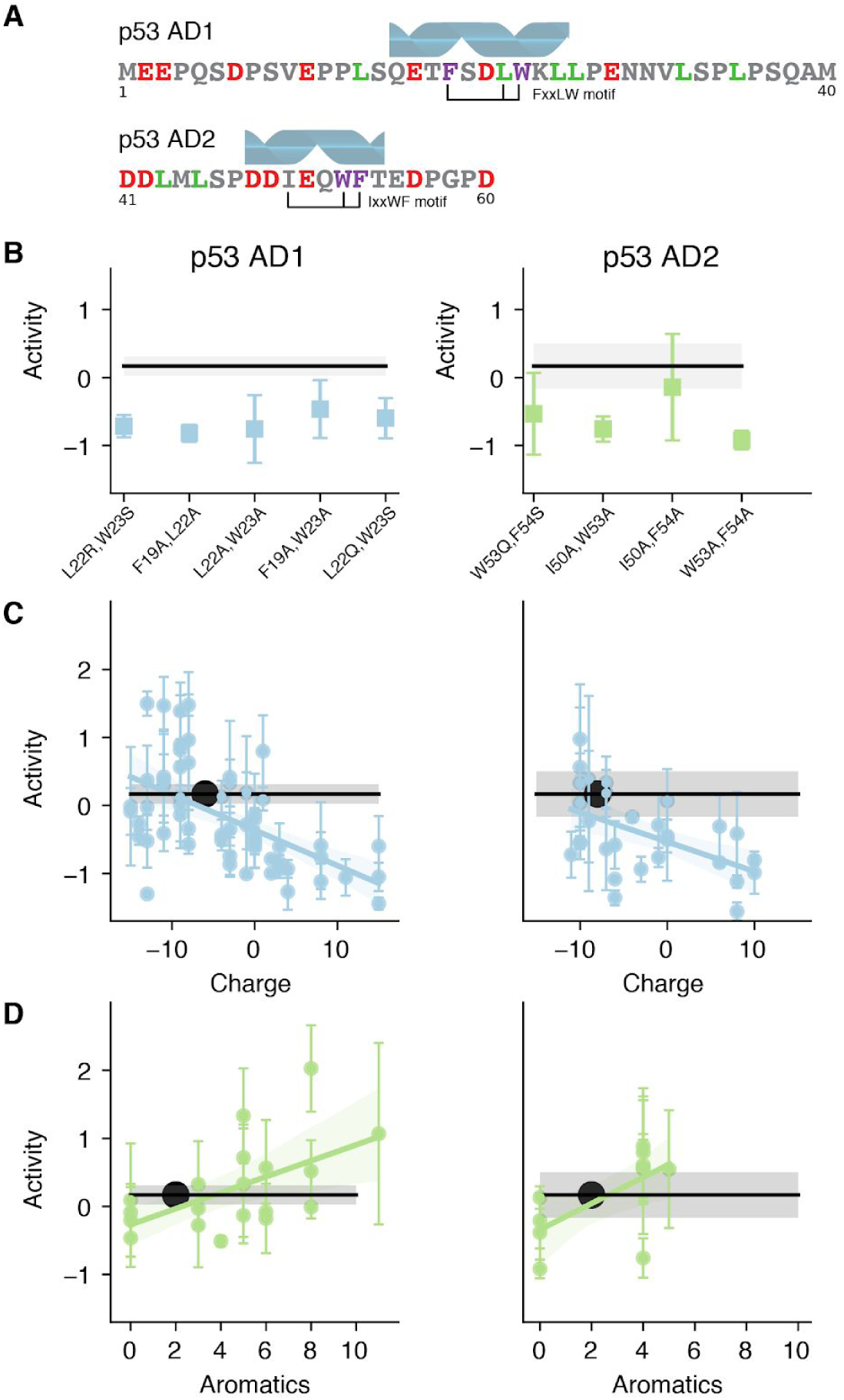
Both activation domains of p53 require acidic residues and hydrophobic motifs for function. A) The motif and alpha helix locations in both p53 ADs. B) Both ADs of p53 contain aromatic and leucine rich motifs that are important for AD function. C) Removing negative charge and/or adding positive charge decreased the activity of both p53 ADs. In p53 AD1, adding acidity (more negative net charge) increases activity in about half of variants. Note that p53 AD2 is only 20 residues, so its net charge per residues is double that of AD1. WT is indicated with a large black dot. WT means and SEMs are shown in all plots with a black line and gray box. Activity is average Z score. A linear regression with a confidence interval summarizes the general dependence on net charge. D) For both ADs, removing aromatic residues generally decreases activity. Adding aromatic residues can increase activity.

**Figure 3–figure supplement 2:**
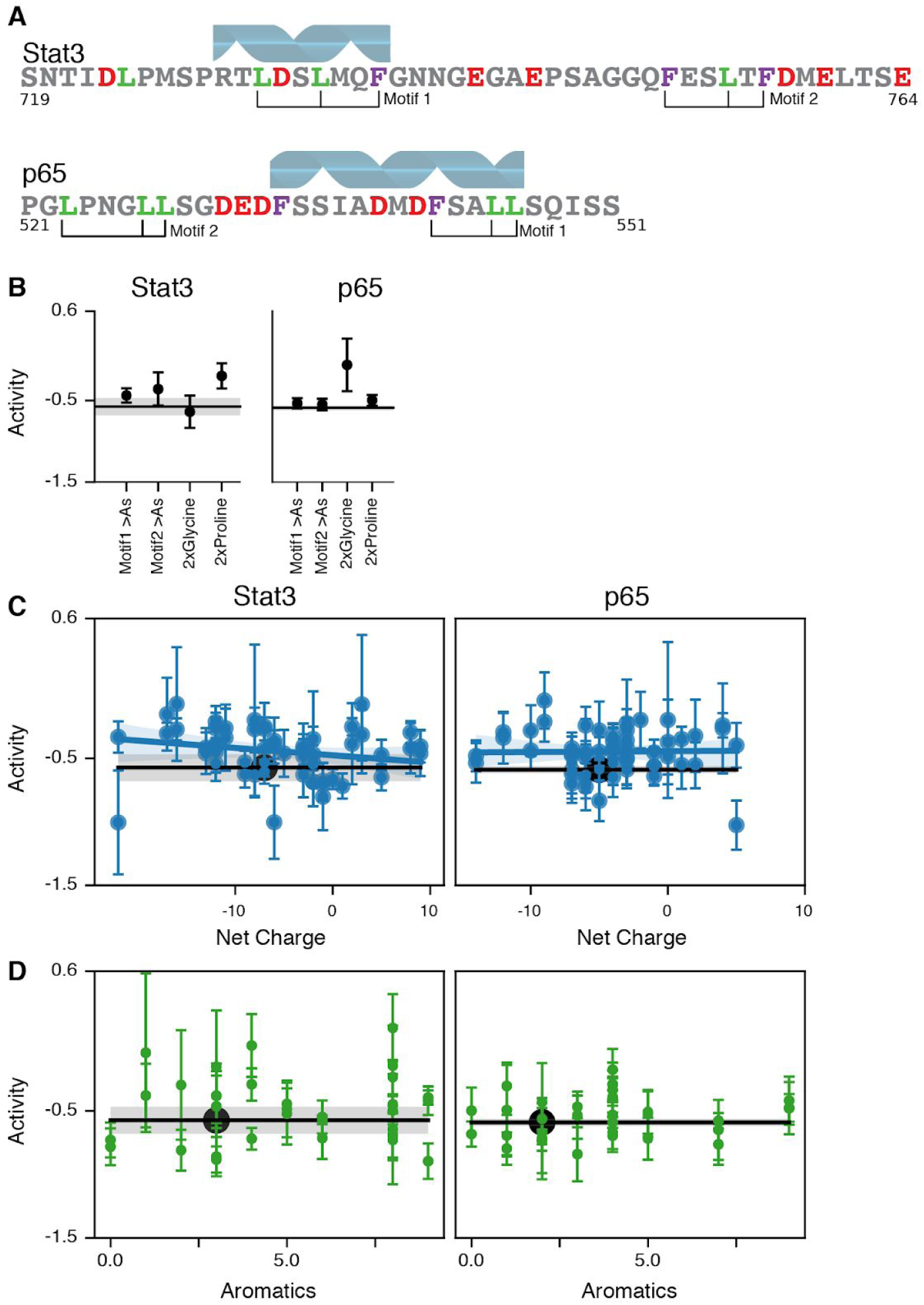
Variants of Stat3 and p65 had very small changes in activity despite large sequence perturbations. A) The motifs and alpha helices of Stat3 and p65. B) The motif mutants and the helix breaking mutants do not cause statistically significant changes in activity. C) The regression suggests that Stat3 activity has a very mild negative correlation with net charge. D) Adding or removing aromatic residues cause small changes in activity. None of the Stat3 or p65 variants had statistically significant changes in activity.

**Figure 3–figure supplement 3:**
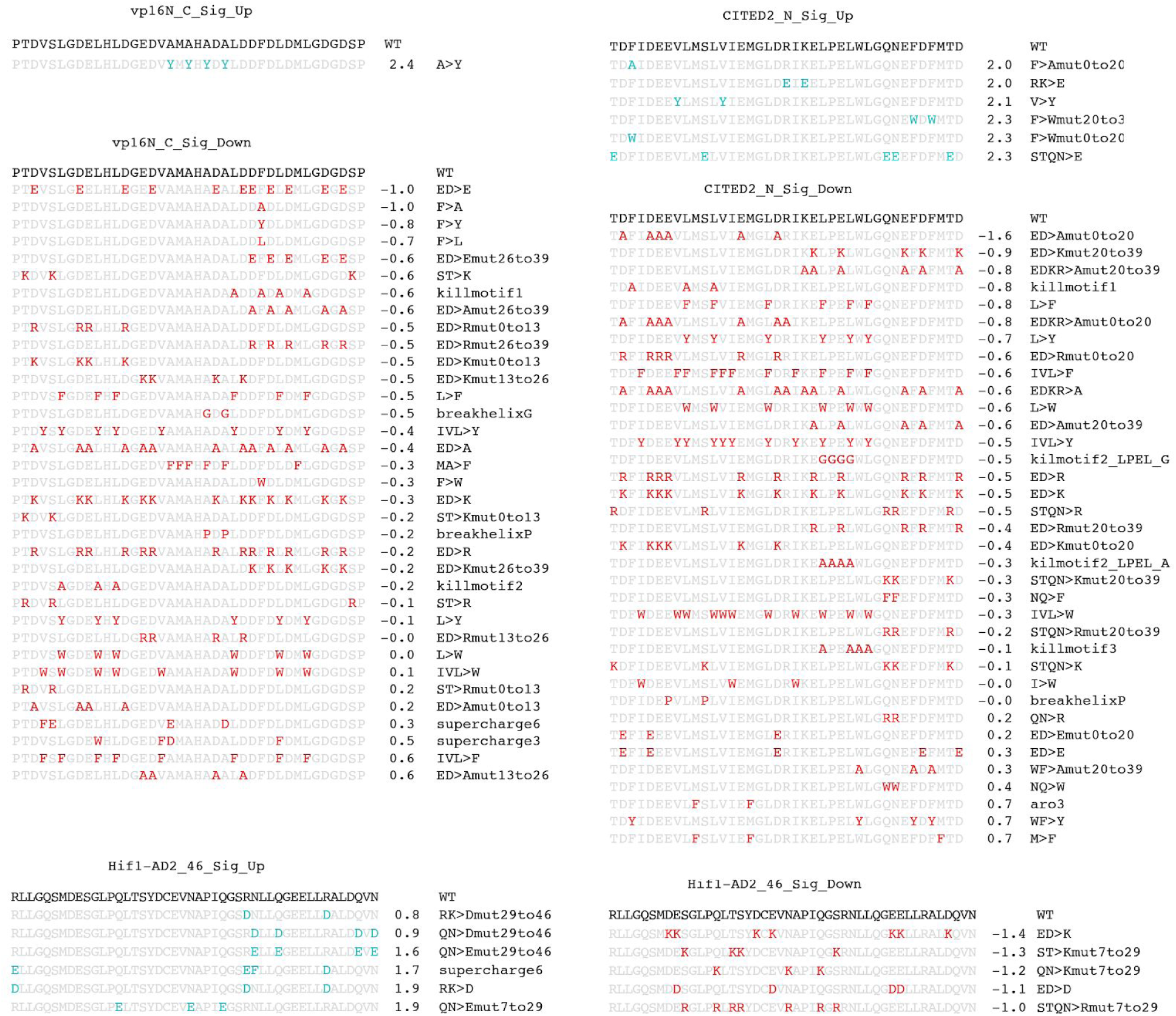
Variants that had a statistically significant increase or decrease activity after correcting for multiple hypothesis testing with 5% FDR. For each AD, variants that are stronger (cyan) or weaker (red) than WT. It is easier to make strong ADs weaker than to make them stronger.

**Figure 3–figure supplement 4:**
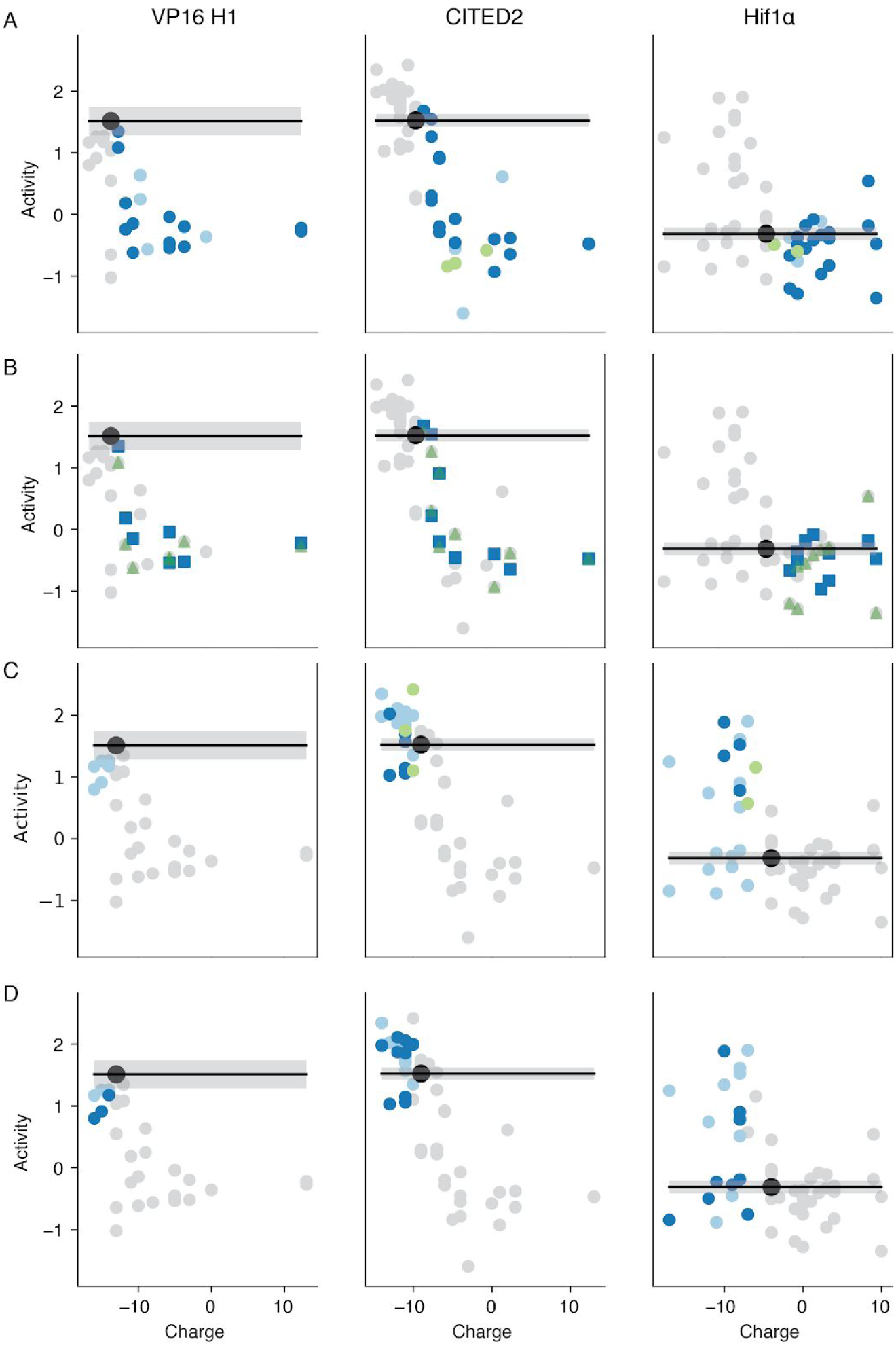
Net charge more efficiently describes the effects of substitutions than amino acid identity. A) Removing acidic residues (negative charges, light blue) has a similar effect as adding basic residues (positive charges, dark blue). B) The identity of added basic residues does not matter. Adding K’s (green) has a similar effect on activity as adding R’s (blue). C) Adding acidic residues (light blue) has similar effects to replacing basic residues with alanine (green) or replacing basic residues with acidic residues (dark blue). In Hif1ɑ, removing positives always increases activity. D) The identities of added acidic residues frequently does not matter. Adding D’s (dark blue) has a similar effect on activity as adding E’s (light blue). In Hif1ɑ, adding E’s is more likely to increase activity than adding D’s.

**Figure 4–figure supplement 1:**
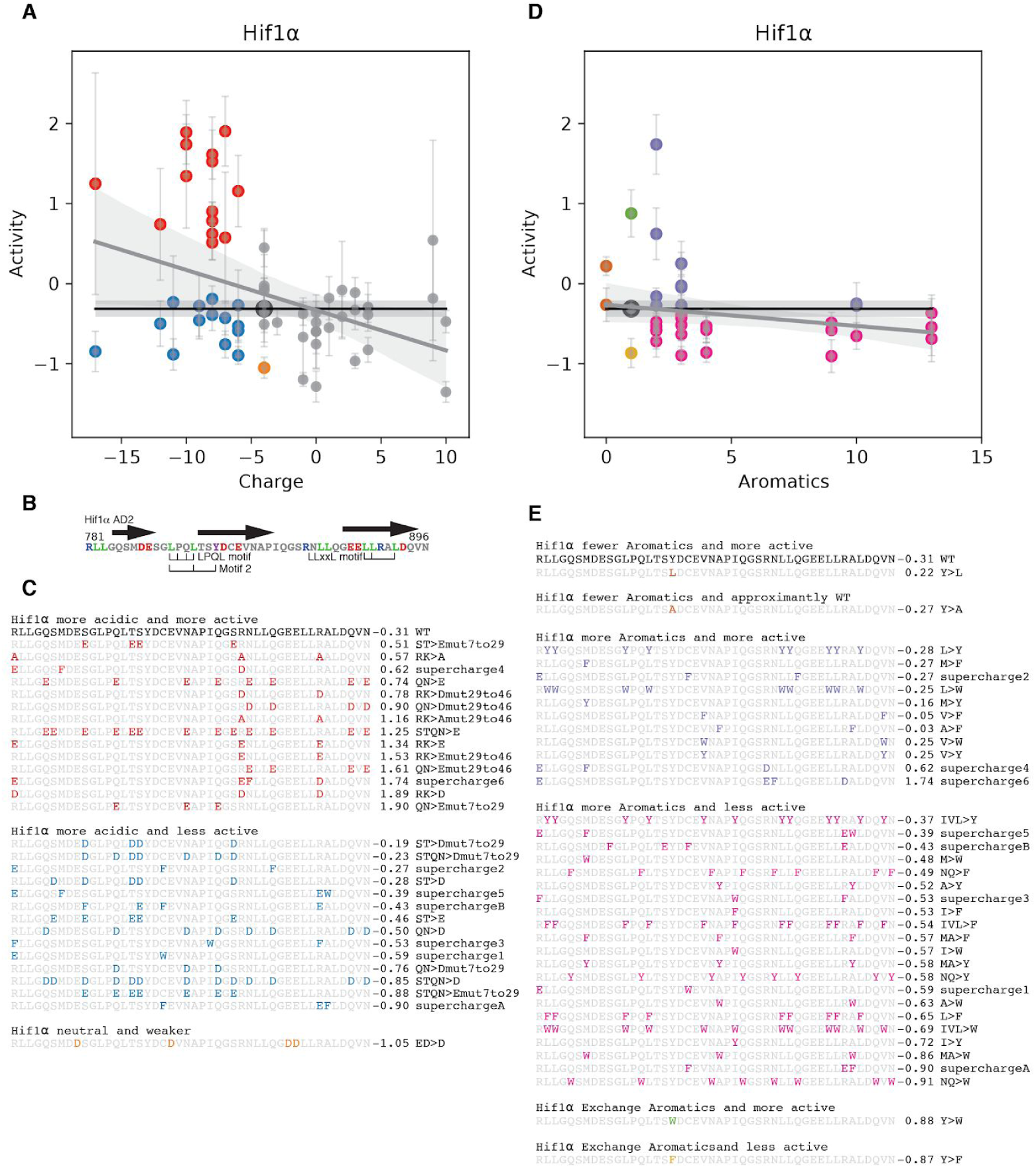
In Hif1ɑ adding negative charges near the motifs is more likely to increase activity A) Variants that added acidity and increased in activity are highlighted in red. The blue variants added acidity and had WT+2*SEM or lower activity. B) The locations of the motifs and alpha helices (arrows). C) The sequences of the red, blue and orange variants from A, in order of increasing activity. The red variants frequently remove the basic residue R820, or add acidic residues near to L795, L812, L813 or L819. The sequences are shown in order of increasing activity. Adding E’s increases activity more often than adding D’s. Replacing E’s with D’s decreased activity (orange), the opposite of the trend observed for VP16 and CITED2. D-E) Adding aromatics to Hif1ɑ generally decreases activity. Replacing the Y with an L causes a small increase in activity. Adding aromatics can increase activity (purple) but more often decreases activity (pink). The two variants that add aromatics and have notably higher activity, supercharge4 and supercharge6, also added acidity. Supercharge6 increased activity in a statistically significant manner, but its similarity to RK>D (panel C) suggests the main effect comes from adding acidity.

**Figure 4–figure supplement 2:**
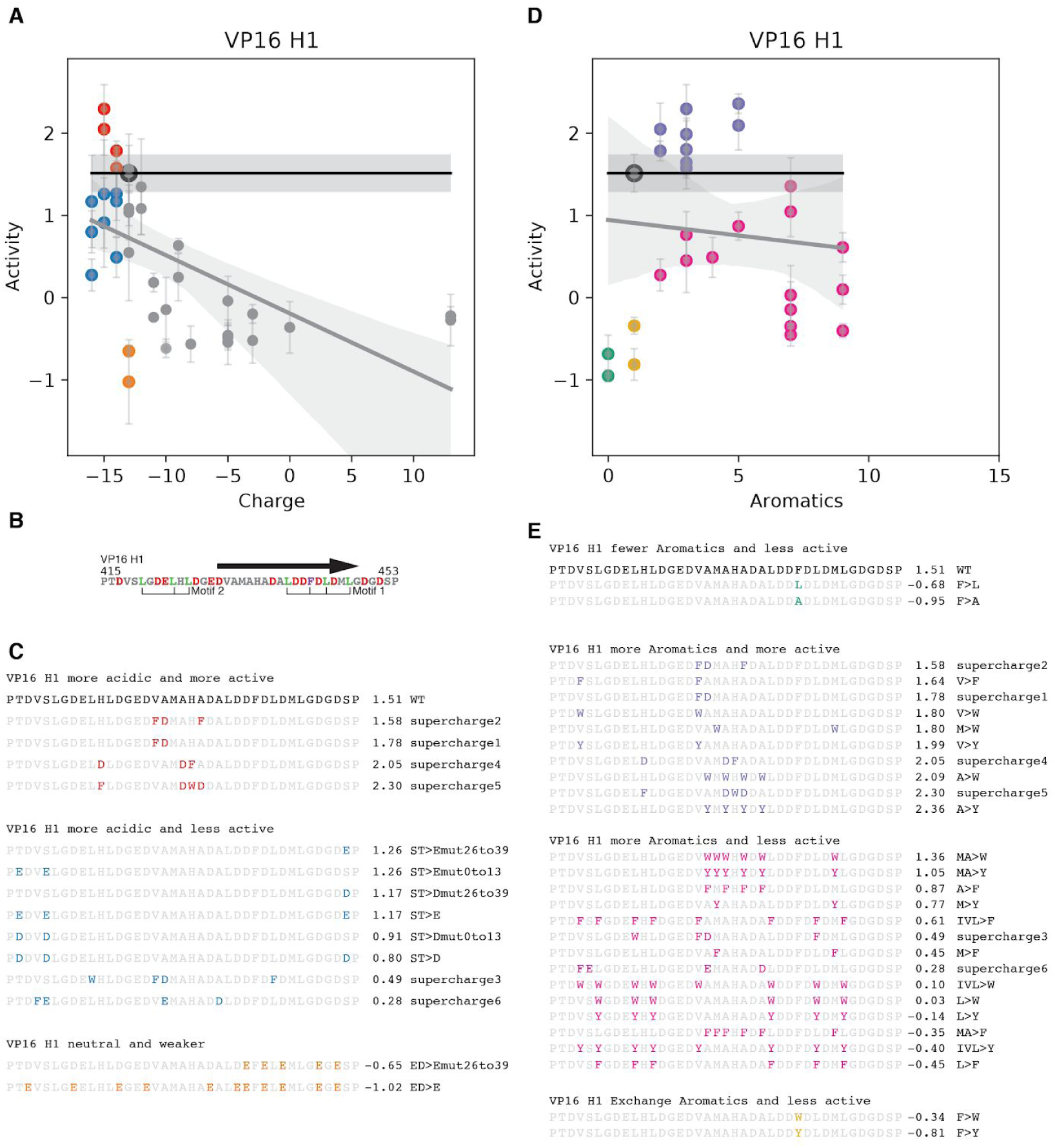
In VP16 adding aromatic residues in the center is more likely to increase activity. A) For VP16, no variants that added acidity increased activity. Some supercharge variants, that added both aromatic and acidic residues, had small increases in activity that were not statistically significant (red). Most variants that added acidity decreased activity (blue). VP16 is saturated for the effect of acidity on activity and this may be because nearly all aromatics and leucines are flanked by acidic residues. B) Motif locations and alpha helix (arrow). C) The locations of variants in A. D>E substitutions in the N terminal region decreased activity (orange). VP16 and CITED2 prefer D over E. D) The one aromatic residue, F442, is critical for activity because all substitutions caused lower activity. Adding 1-4 aromatics can increase activity (purple), but often does not (pink). E) The purple variants tended to add aromatics in the center of the AD, N-terminal to F442. Adding more aromatic residues or adding them C-terminal to F442 tended to decrease activity. The only variant that increased activity in a statistically significant manner was A>Y.

**Figure 4–figure supplement 3:**
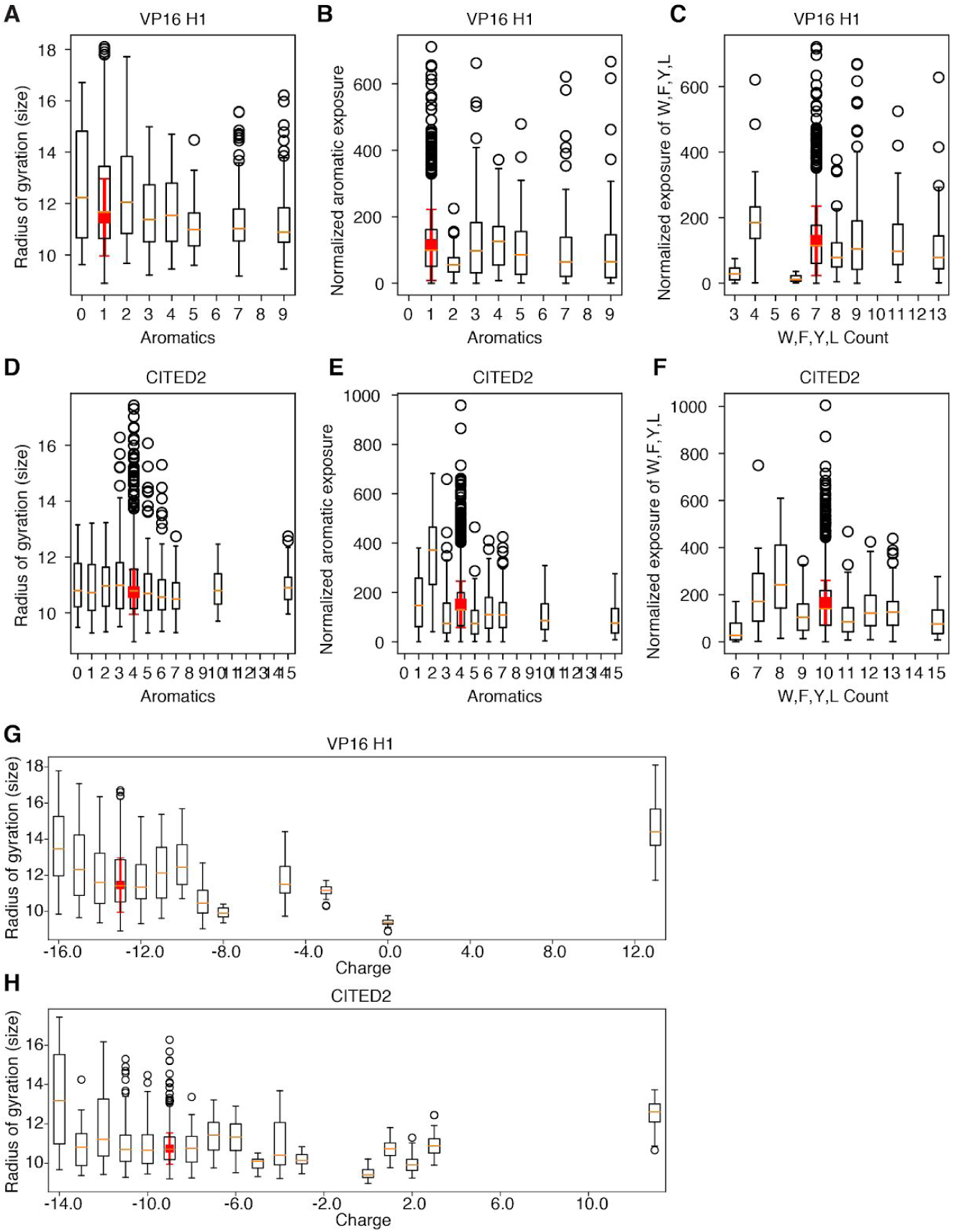
Summary of all atom Monte Carlo simulations of VP16 and CITED2 variants. These simulations are well-poised to capture the conformational ensemble of and residual structure in disordered proteins. Simulations of the Yeast AD, Gcn4, helped us develop the Acidic Exposure Model (Staller et al., 2018). For each variant, we ran 10 simulations starting in a helix and 10 simulations starting in a random coil. Here, each of the 20 simulations for each variant is included separately. For VP16 (A) and CITED2 (B) variants with more aromatic residues tended to have a smaller radius of gyration, a parameter that captures the diameter of the conformational ensemble. In both ADs, adding aromatic residues leads to smaller radius of gyration, consistent with chain collapse. (C-F) For each simulation, we computed the exposure of all aromatic residues (C-D) or W,F,Y&L residues (E-F) (See Methods) and normalized this total exposure by the number of aromatics or W,F,Y&L residues in each sequence. For VP16, adding 2-3 aromatics leads to a slight increased normalized exposure. For CITED2, adding aromatics generally does not increase normalized exposure. (G-H) Adding acidic residues frequently increased the radius of gyration, indicating a more expanded ensemble of conformations.

**Figure 5–figure supplement 1:**
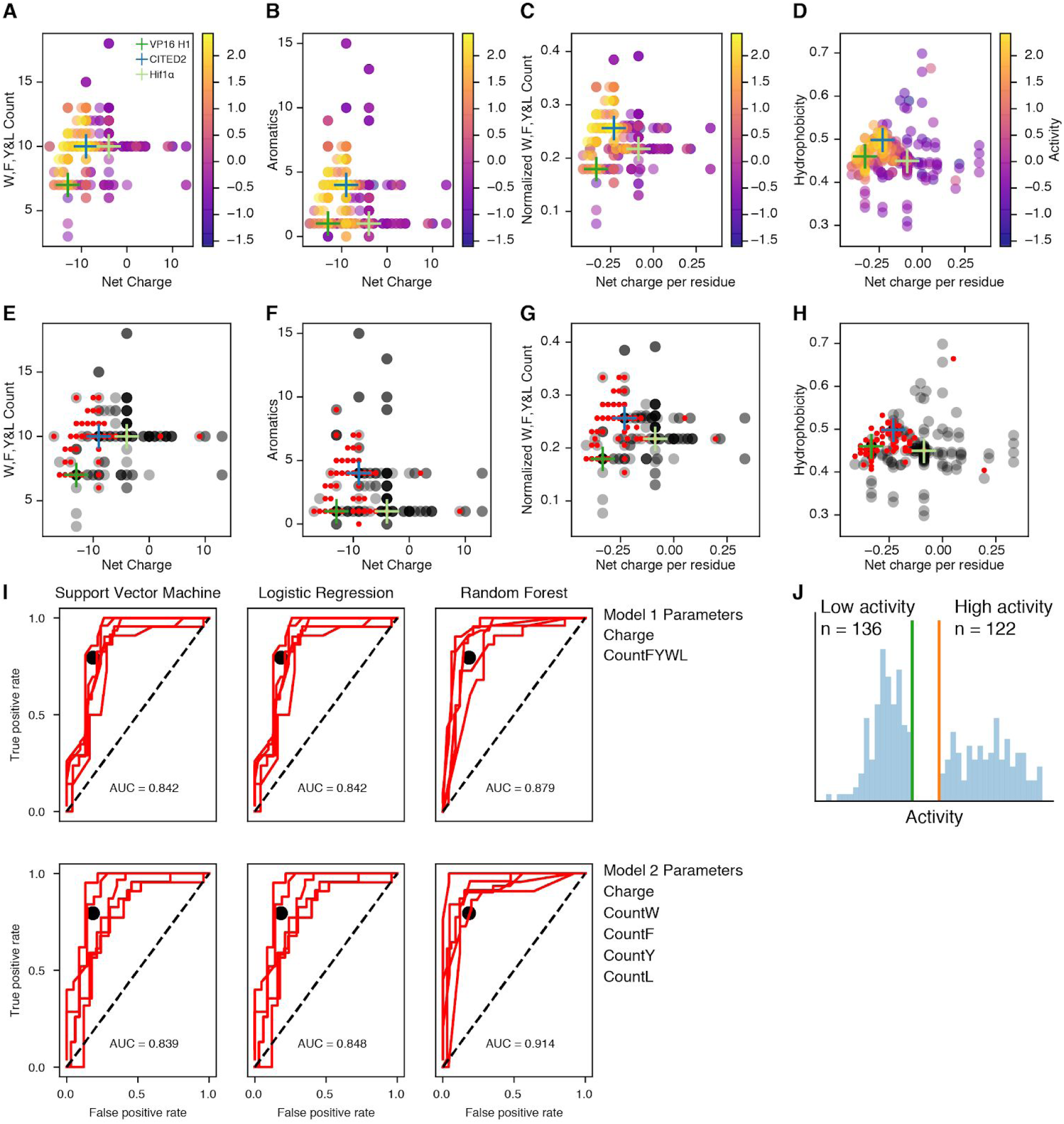
Net charge and W,F,Y&L count can separate high and low activity variants. A) Repeat of Figure. 5A. The location of each point encodes its properties. Color indicates activity. B) Counting only aromatics does not separate high and low activity variants as well as counting W,F,Y&L residues. C) Normalizing the counts and charge by AD length preserves the separation between high and low activity variants. D) Using Kyte-Doolittle hydrophobicity(Kyte and Doolittle, 1982) instead of counting W,F,Y&L also separates high and low activity variants. WT ADs are indicated with crosses. E-H) Replotting A-D with binarized data. Many points on this grid are occupied by both high (red) and low (gray) activity variants. We split the data into high (Activity >0.5, N=125, red) and low (Activity <=0.5, N = 177, gray) activity variants and replotted them. When multiple low points overlap they appear black. Red points are on top of gary points to emphasize the overlap. I) We deployed Support Vector Machines, Logistic Regression and Random Forest classifiers. We included all VP16, Hif1ɑ and CITED2 variants except for the shuffle variants, because these composition-based classification models cannot distinguish the shuffle variants from WT. Receiver operator characteristic curves for 5-fold cross validation. The average area under the curve (AUC) of the 5 test sets is shown. For comparison, the performance of the proteome-wide AD predictor described in Figure 5C is plotted as a black dot. J) There are 122 data points in the High Activity Set (Activity >0.5) and 136 in the Low Activity Set (Activity <0, see Methods).

**Figure 5–figure supplement 2:**
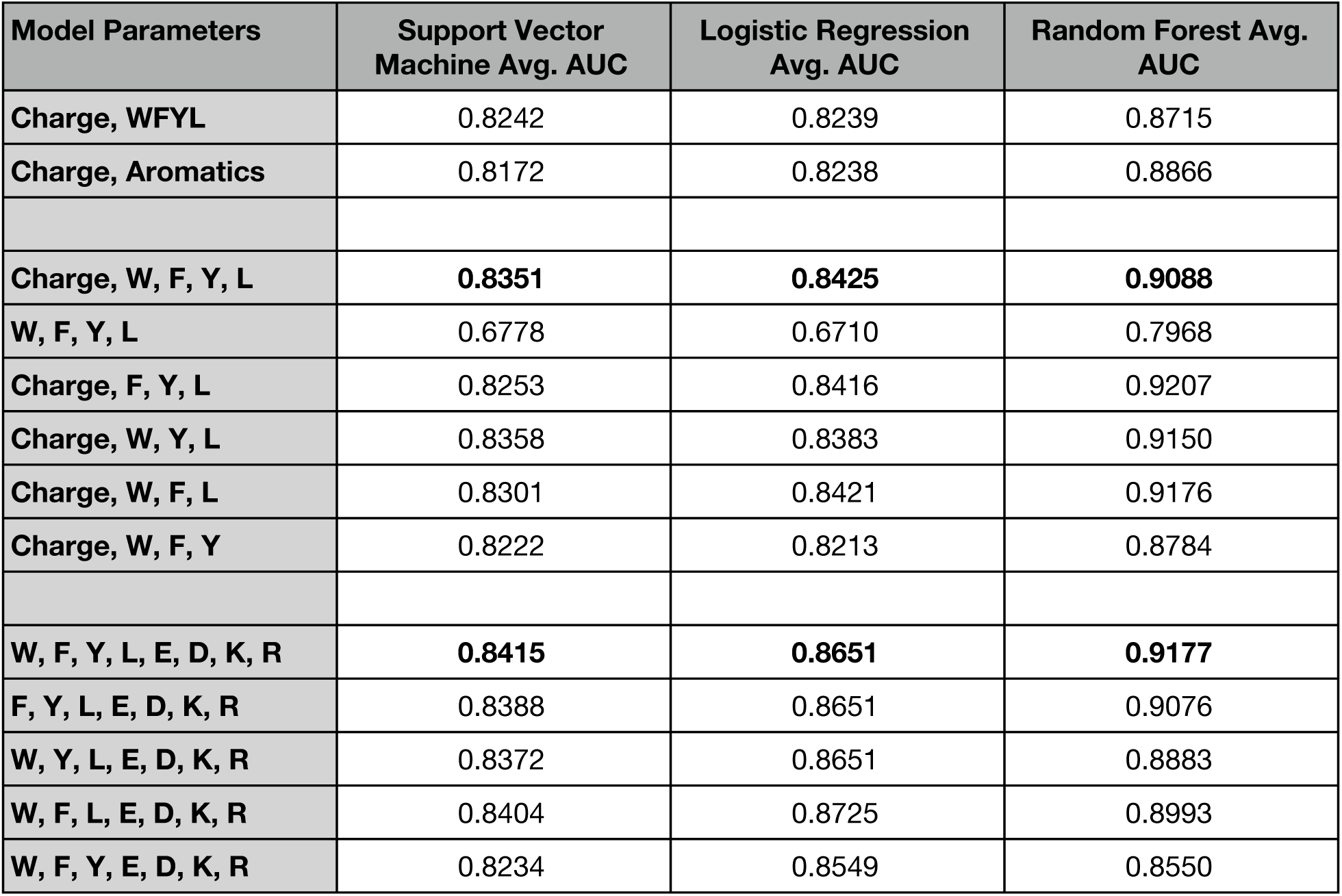
Comparison of machine learning models with different parameter sets. For each parameter set, the average AUCs from 5 fold cross validation are shown. Net Charge is the parameter that, when removed, caused the largest drop in model performance. Removing leucine residues caused a larger drop in model performance than removing any individual aromatic residue.

**Figure 6–figure supplement 1:**
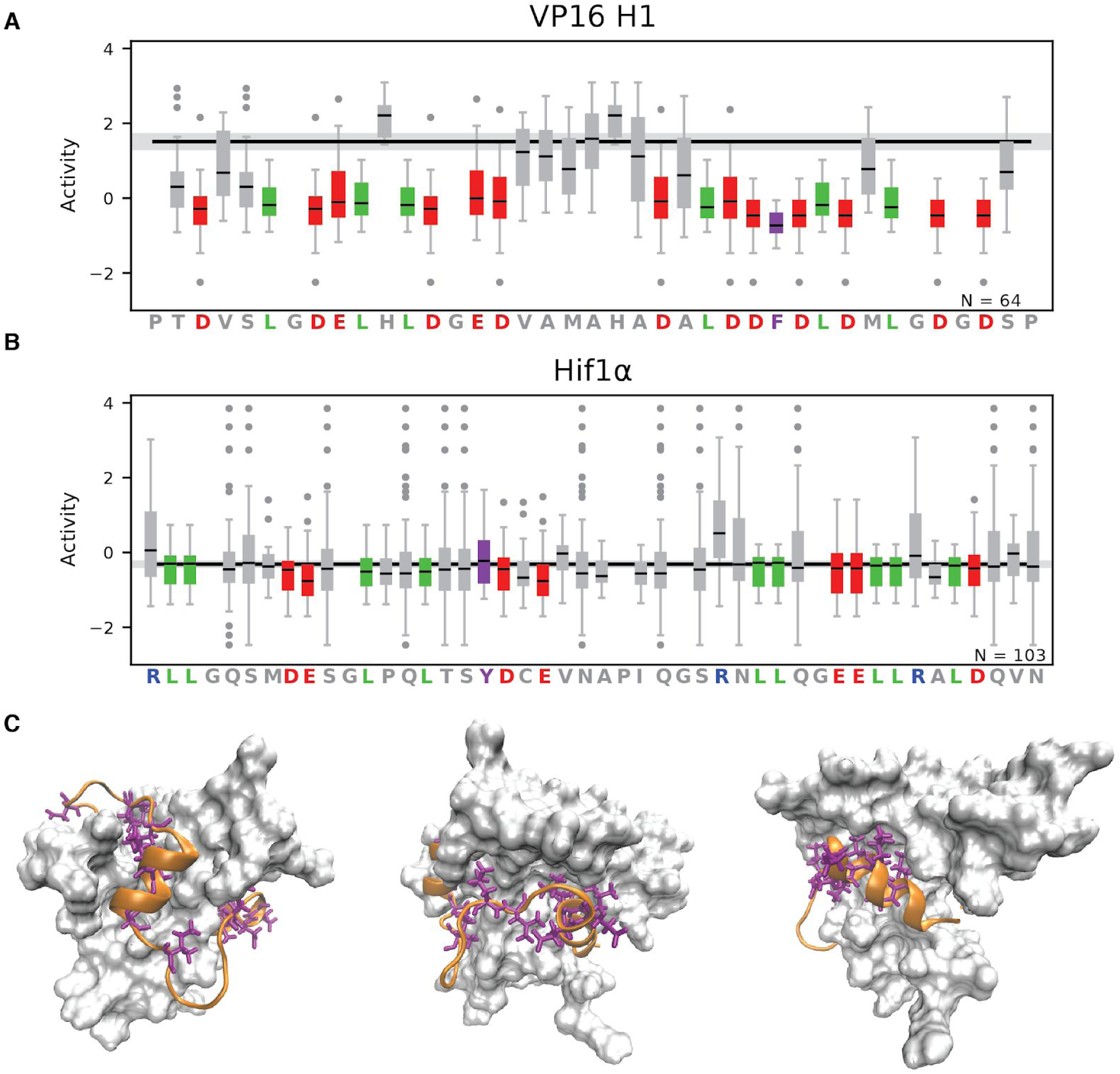
Leucines and acidic residues contribute to activity of VP16 and Hif1ɑ. A-B) For each position, the activities of all variants that introduced a substitution at that position are summarized. The 4 biological replicates are included separately. Most variants changed multiple positions. Each residue was mutated a different number of times (1-22 variants, 4-88 measurements) and mutated to different amino acids. A) In VP16, F442 is critical for activity. The leucine and acidic residues also contribute to activity. B) For Hif1ɑ, WT activity was low, reducing our ability to resolve decreases in activity. Leucines and acidic residues have low average activities. C) In the structure of the Hif1ɑ-TAZ1 interaction, positions with the lowest average activities (purple) in Hif1ɑ (orange) point towards the surface of the TAZ1 (white). Three rotations of the same snapshot are shown. The mutagenesis can detect positions that contact the coactivator.

**Figure 6–figure supplement 2:**
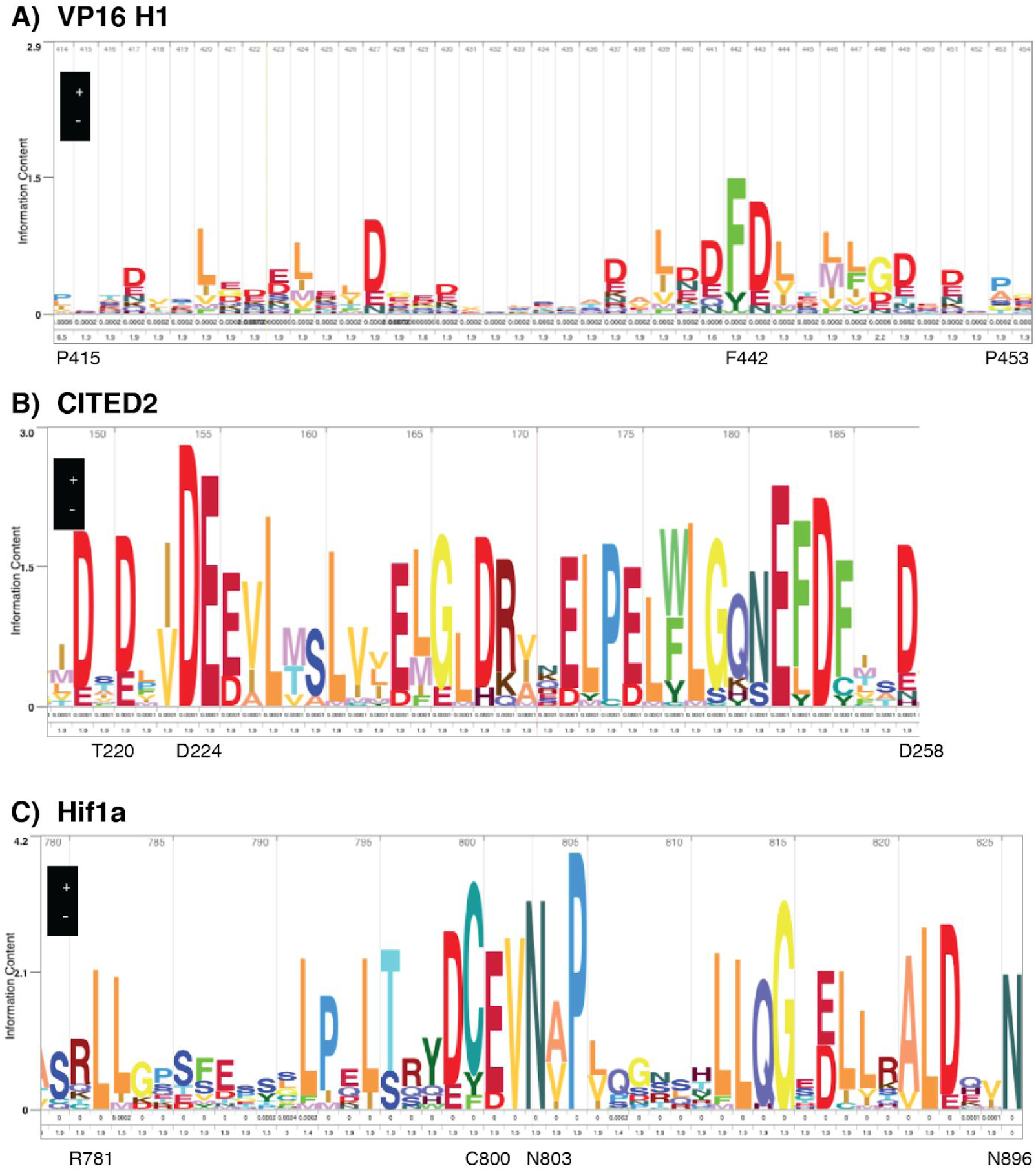
The HMM logos from HMMER for each AD. For each AD, we ran HMMER for 3 rounds, at which point VP16 and CITED2 had converged and Hif1ɑ had >126K sequences. We show the HMMER generated logo for the Hidden Markov Model (HMM) model for each AD region. The height of each letter encodes the information content of residue, a reasonable proxy for conservation. A) For VP16, F442, the leucine and aspartic acid residues are the positions with the highest information. In our data, these positions make large contributions to activity. B) For CITED2, D224 is the position in the HMM with the most information. In our data this residue contributes to activity. In addition, the leucine and acidic residues that contribute to activity in our data have high information content. C) In Hif1ɑ, C800 is important for sensing hypoxia and N803 is hydroxylated under normoxia, a modification that interferes with binding to p300 (Lando et al., 2002; Yasinska and Sumbayev, 2003). The leucine and acidic residues that contribute to activity in our data have moderate to high information content.

**Figure 6–figure supplement 3:**
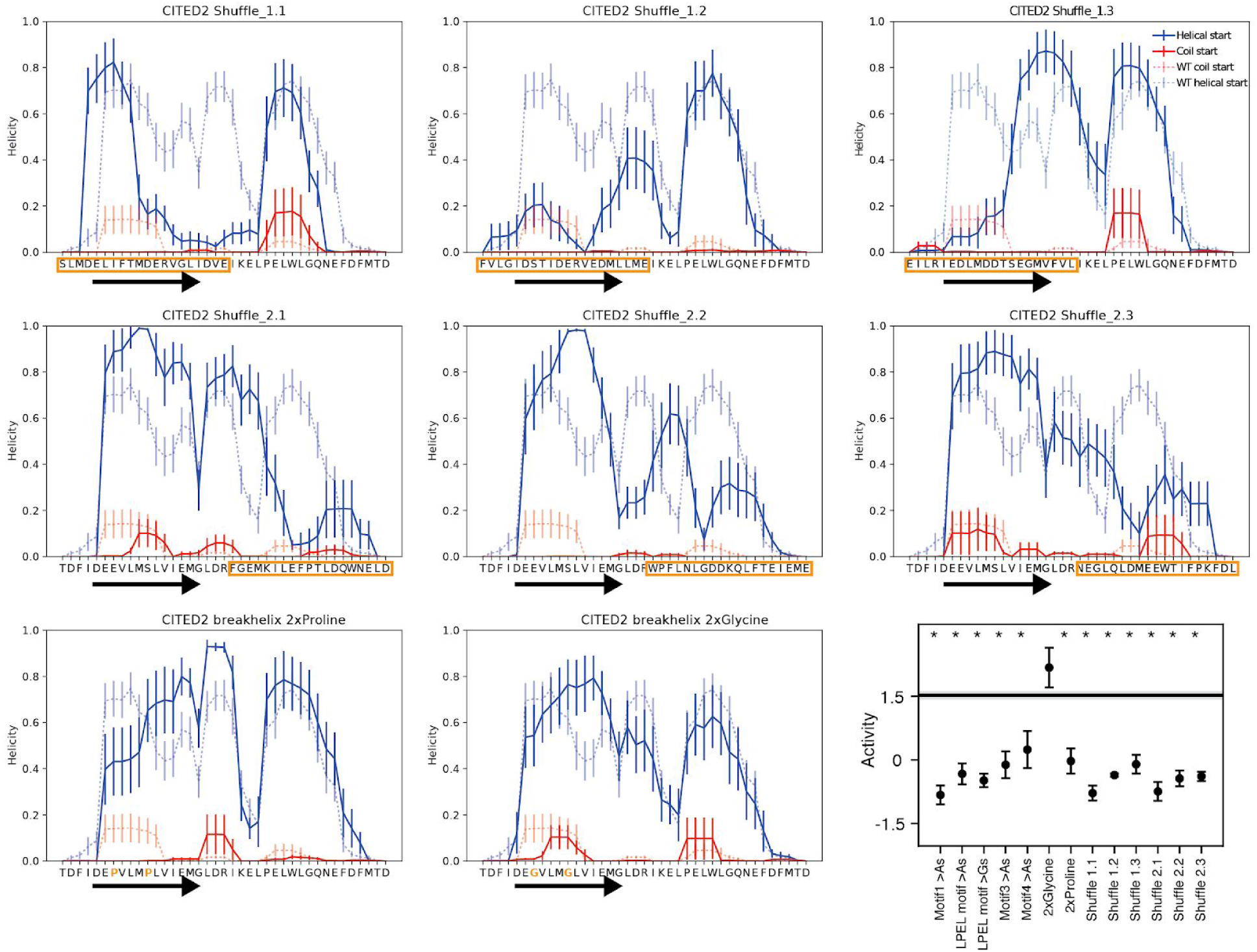
Mutations designed to disrupt alpha helix formation in CITED frequently reduce helicity in the all-atom Monte Carlo simulations. For each variant, we ran 10 simulations starting as a helix (solid blue) and 10 starting as a random coil (solid red). The arrow indicates the alpha helix observed in the NMR interaction structure with TAZ1. Not all simulations converged between these starting conditions, so it is only appropriate to compare each starting condition to its matched WT simulations (dashed lines). For example Shuffle variant 1.1 shuffled the N terminal region (orange box). In the random coil set, helicity in the NMR helix region disappeared (compare solid red line to dashed red line). In the helix set, helicity in the center was greatly reduced. We conclude that this variant reduced helicity. The 2xProline variant, in the random coil set, eliminated helicity over the NMR helix and increased helicity in the center. The 2xGlycine variant retained some helicity in the NMR region. The last panel shows the activities of these variants and motif disrupting mutants for comparison. Shuffling any region of this AD decreased activity as much as removing a motif, indicating that the arrangement (sequence) of residues is important for function.

**Figure 6–figure supplement 4:**
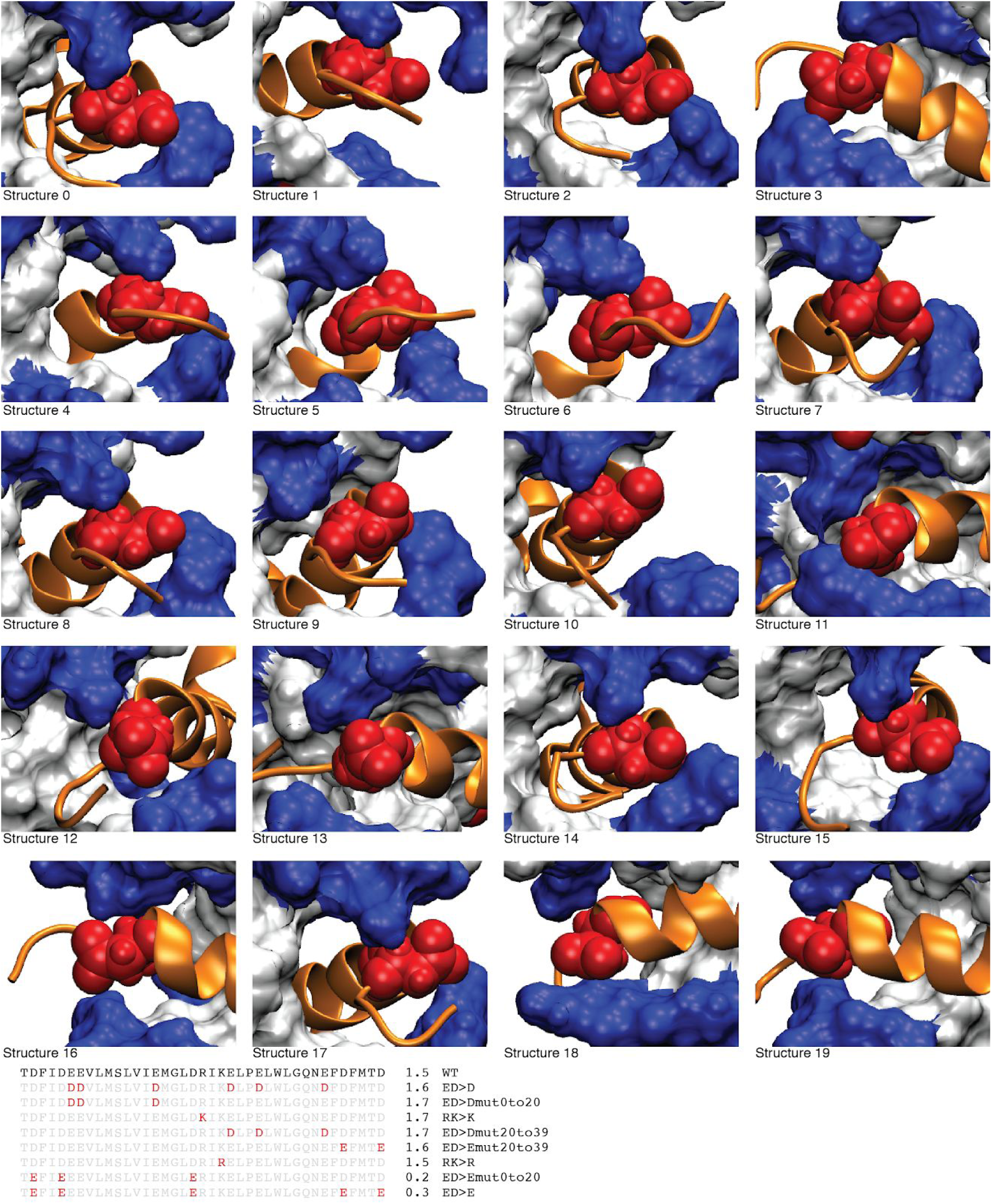
Snapshots of CITED2 bound to the TAZ1 domain of CBP. In 18/20 structures (1R8U), D224 (red) of CITED2 (orange) sits between the narrow, positively charged rims (blue) of the binding canyon on TAZ1 (white). CITED2 D224 is closest to R439 (upper blue) and K365 (lower blue) of TAZ1. The canyon is widest in structures 1 and 10, and D224 interacts primarily with R439. Structure 16 is shown in Figure 6C. Most variants that exchanged charged residues had nearly WT activity, but two variants with the D221E, D224E and D238 substitutions had decreased activity. D221 and D238 are not sterically constrained.

**Figure 6–figure supplement 5:**
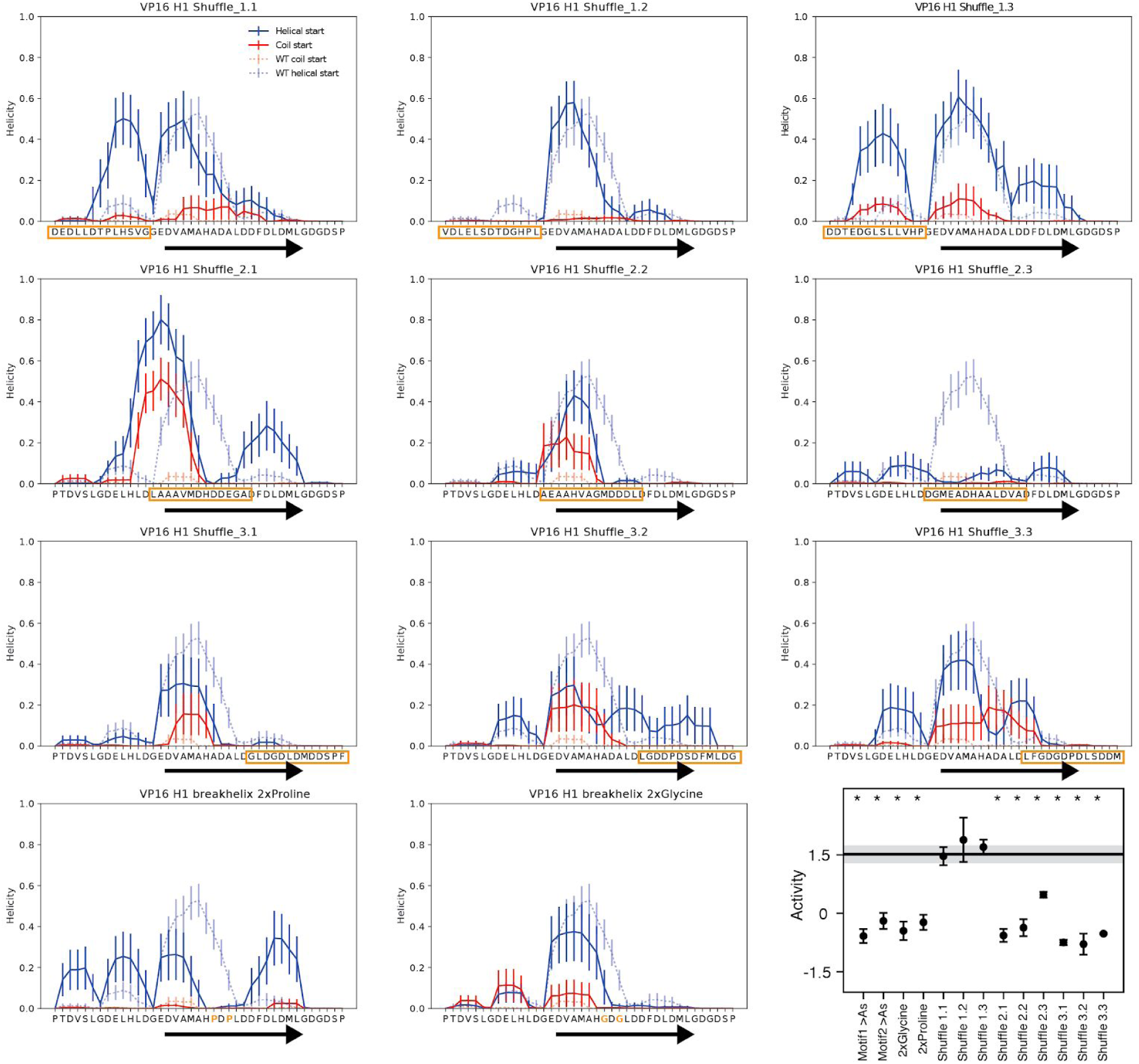
Mutations designed to disrupt the alpha helix of VP16 frequently reduce helicity in simulations. For each variant, we ran 10 simulations starting as a helix (solid blue) and 10 starting as a random coil (solid red). Not all simulations converged between these starting conditions, so it is only appropriate to compare each starting condition to its matched WT simulations (dashed lines). Shuffling the sequence of region 1 did not disrupt the helix and did not decrease AD activity. Shuffle variants 2.1, 2.2 and 3.1 moved the helix toward the N terminus. Variant 2.3 decreased helicity. Variants 3.2 and 3.3 perturbed helicity. The 2xProline and 2xGlycine variants disrupted helicity near the substitutions. The inset plot shows the activities of all the variants and motif disrupting variants for comparison. Variants that shuffled regions that overlapped the known helix (black arrow) reduced activity. Shuffling Region 1 did not reduce activity, but mutating the leucines in this region to alanines did reduce activity. This region appears to have fewer constraints on the arrangement of residues, and may exhibit fuzzy binding to a coactivator.

**Figure 6–figure supplement 6:**
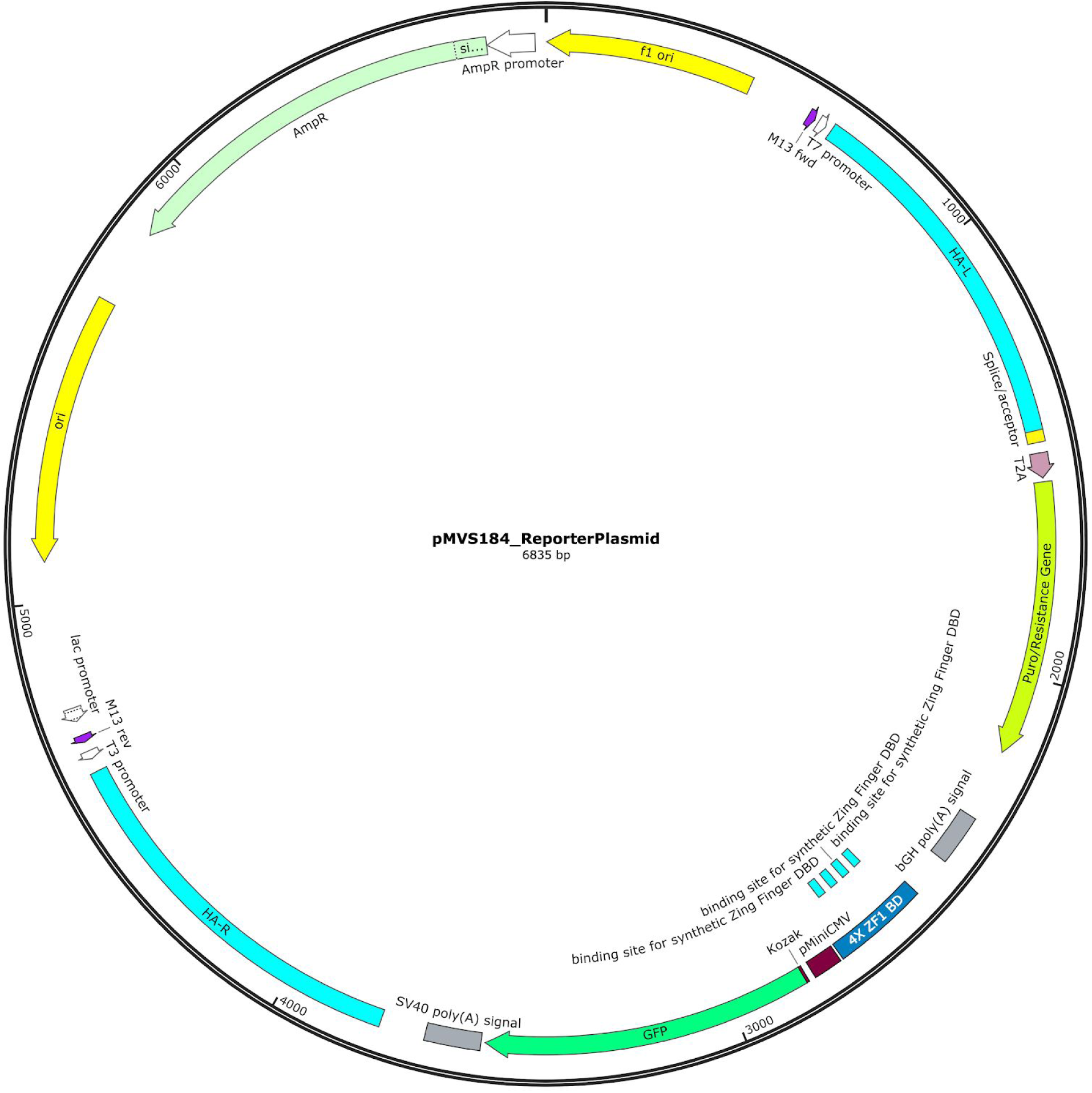
The GFP reporter, pMVS184. This plasmid has four binding sites for the synthetic DBD and is integrated at the AAVS1 locus. The synthetic zinc finger DBD and cognate TF binding sites were designed and validated by Minhee Park and Ahmed Khalil (Park et al., 2019). The reporter plasmid (pMVS184) contains upstream homology to AAVS1 (804 bp), a splice acceptor, a T2A signal, puromycin resistance, the bGH poly(A) signal, four binding sites for the synthetic zinc finger DBD, the pMiniCMV minimal promoter, GFP, the SV40 3’ UTR and downstream homology to AAVS1 (837 bp). The plasmid sequence is in Supplemental Dataset 6. The plasmid will be available from Addgene.

**Figure 6–figure supplement 7:**
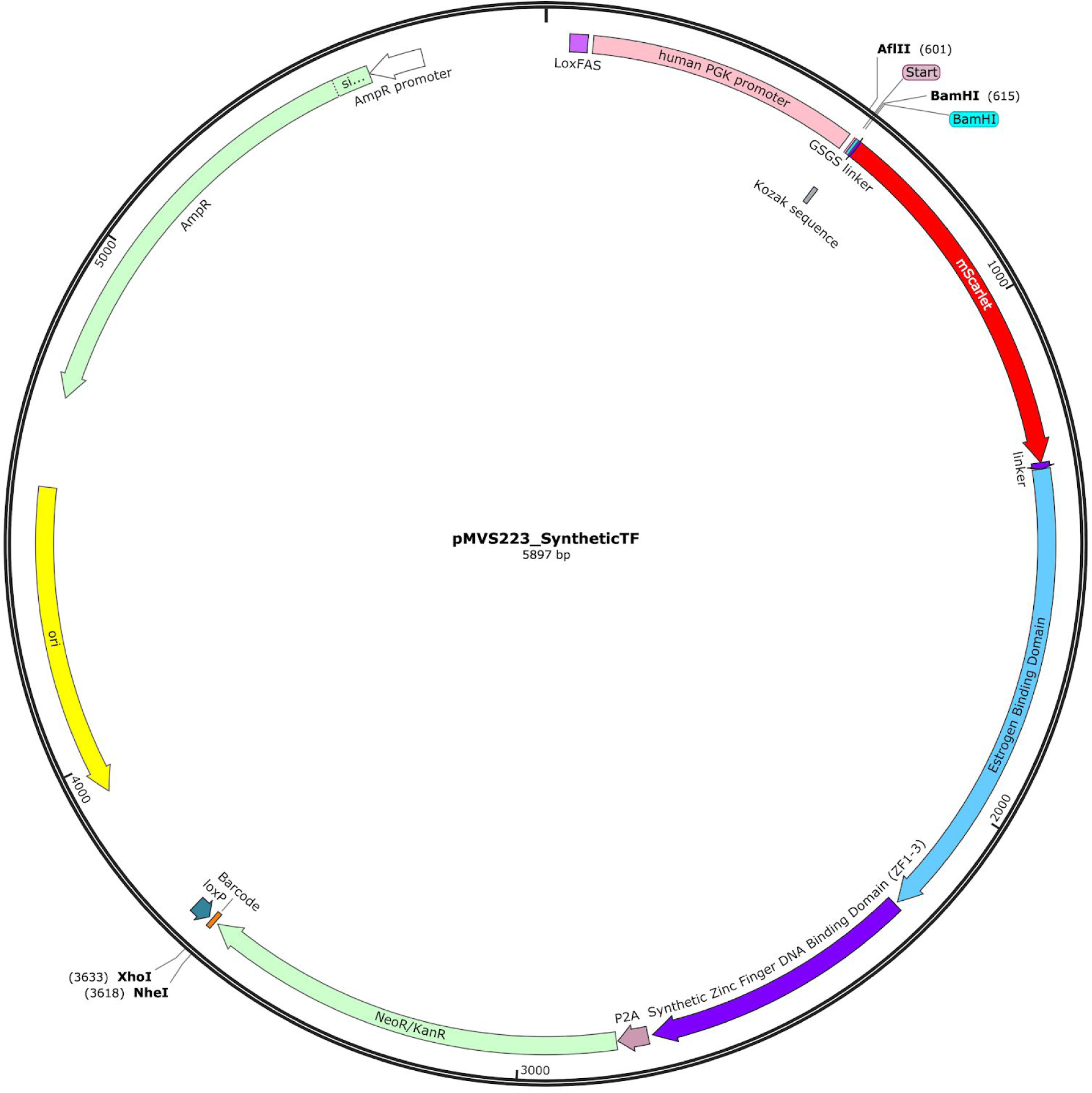
Map of the synthetic TF plasmid, pMVS223. The synthetic TF contains a loxFAS site, the human PGK promoter, a multiple cloning site for inserting AD variants, an mScarlet red fluorescent protein, an estrogen response domain, a synthetic zinc finger DNA binding domain (DBD), a P2A cleavage sequence in frame with a neomycin resistance gene, a stop codon, a barcode sequence in the 3’ UTR, and a loxP site. The landing pad adds a WPRE sequence to the 3’ UTR of the final transcript (Maricque et al., 2018). In the final library, the AD variants are located between the ATG START and the BamHI site, and the barcodes are between the NheI and XhoI sites. The plasmid sequence is in Supplemental Dataset 6. The plasmid will be available from Addgene.

## References

1. Andresen C, Helander S, Lemak A, Farès C, Csizmok V, Carlsson J, Penn LZ, Forman-Kay JD, Arrowsmith CH, Lundström P, Sunnerhagen M. 2012. Transient structure and dynamics in the disordered c-Myc transactivation domain affect Bin1 binding. Nucleic Acids Res 40:6353–6366. doi:10.1093/nar/gks263

2. Berlow RB, Dyson HJ, Wright PE. 2017. Hypersensitive termination of the hypoxic response by a disordered protein switch. Nature 543:447–451. doi:10.1038/nature21705

3. Boija A, Klein IA, Sabari BR, Dall’Agnese A, Coffey EL, Zamudio AV, Li CH, Shrinivas K, Manteiga JC, Hannett NM, Abraham BJ, Afeyan LK, Guo YE, Rimel JK, Fant CB, Schuijers J, Lee TI, Taatjes DJ, Young RA. 2018. Transcription Factors Activate Genes through the Phase-Separation Capacity of Their Activation Domains. Cell 175:1842–1855.e16. doi:10.1016/j.cell.2018.10.042

4. Brzovic PS, Heikaus CC, Kisselev L, Vernon R, Herbig E, Pacheco D, Warfield L, Littlefield P, Baker D, Klevit RE, Hahn S. 2011. The acidic transcription activator Gcn4 binds the mediator subunit Gal11/Med15 using a simple protein interface forming a fuzzy complex. Mol Cell 44:942–953. doi:10.1016/j.molcel.2011.11.008

5. Chang J, Kim DH, Lee SW, Choi KY, Sung YC. 1995. Transactivation ability of p53 transcriptional activation domain is directly related to the binding affinity to TATA-binding protein. J Biol Chem 270:25014–25019. doi:10.1074/jbc.270.42.25014

6. Chen S, Wang Q-L, Xu S, Liu I, Li LY, Wang Y, Zack DJ. 2002. Functional analysis of cone–rod homeobox (CRX) mutations associated with retinal dystrophy. Hum Mol Genet 11:873–884. doi:10.1093/hmg/11.8.873

7. Choi Y, Asada S, Uesugi M. 2000. Divergent hTAFII31-binding motifs hidden in activation domains. J Biol Chem 275:15912–15916. doi:10.1074/jbc.275.21.15912

8. Chong S, Dugast-Darzacq C, Liu Z, Dong P, Dailey GM, Cattoglio C, Heckert A, Banala S, Lavis L, Darzacq X, Tjian R. 2018. Imaging dynamic and selective low-complexity domain interactions that control gene transcription. Science 361. doi:10.1126/science.aar2555

9. Cress WD, Triezenberg SJ. 1991. Critical structural elements of the VP16 transcriptional activation domain. Science 251:87–90.

10. Diss G, Lehner B. 2018. The genetic landscape of a physical interaction. Elife 7. doi:10.7554/eLife.32472

11. Drysdale CM, Duenas E, Jackson BM, Reusser U, Braus GH, Hinnebusch AG. 1995. The transcriptional activator GCN4 contains multiple activation domains that are critically dependent on hydrophobic amino acids. Mol Cell Biol 15:1220–1233.

12. Dyson HJ, Wright PE. 2016. Role of Intrinsic Protein Disorder in the Function and Interactions of the Transcriptional Coactivators CREB-binding Protein (CBP) and p300. J Biol Chem 291:6714–6722. doi:10.1074/jbc.R115.692020

13. El-Gebali S, Mistry J, Bateman A, Eddy SR, Luciani A, Potter SC, Qureshi M, Richardson LJ, Salazar GA, Smart A, Sonnhammer ELL, Hirsh L, Paladin L, Piovesan D, Tosatto SCE, Finn RD. 2019. The Pfam protein families database in 2019. Nucleic Acids Res 47:D427–D432. doi:10.1093/nar/gky995

14. Erijman A, Kozlowski L, Sohrabi-Jahromi S, Fishburn J, Warfield L, Schreiber J, Noble WS, Söding J, Hahn S. 2020. A High-Throughput Screen for Transcription Activation Domains Reveals Their Sequence Features and Permits Prediction by Deep Learning. Mol Cell 78:890–902.e6. doi:10.1016/j.molcel.2020.04.020

15. Ferreira ME, Hermann S, Prochasson P, Workman JL, Berndt KD, Wright APH. 2005. Mechanism of transcription factor recruitment by acidic activators. J Biol Chem 280:21779–21784. doi:10.1074/jbc.M502627200

16. Finn RD, Coggill P, Eberhardt RY, Eddy SR, Mistry J, Mitchell AL, Potter SC, Punta M, Qureshi M, Sangrador-Vegas A, Salazar GA, Tate J, Bateman A. 2016. The Pfam protein families database: towards a more sustainable future. Nucleic Acids Res 44:D279–85. doi:10.1093/nar/gkv1344

17. Freedman SJ, Sun Z-YJ, Kung AL, France DS, Wagner G, Eck MJ. 2003. Structural basis for negative regulation of hypoxia-inducible factor-1α by CITED2. Nat Struct Mol Biol 10:504–512. doi:10.1038/nsb936

18. Hawkins JA, Jones SK Jr, Finkelstein IJ, Press WH. 2018. Indel-correcting DNA barcodes for high-throughput sequencing. Proc Natl Acad Sci U S A 115:E6217–E6226. doi:10.1073/pnas.1802640115

19. Hermann S, Berndt KD, Wright AP. 2001. How transcriptional activators bind target proteins. J Biol Chem 276:40127–40132. doi:10.1074/jbc.M103793200

20. Holehouse AS, Das RK, Ahad JN, Richardson MOG, Pappu RV. 2017. CIDER: Resources to Analyze Sequence-Ensemble Relationships of Intrinsically Disordered Proteins. Biophys J 112:16–21. doi:10.1016/j.bpj.2016.11.3200

21. Hope IA, Struhl K. 1986. Functional dissection of a eukaryotic transcriptional activator protein, GCN4 of yeast. Cell 46:885–894.

22. Humphrey W, Dalke A, Schulten K. 1996. VMD: visual molecular dynamics. J Mol Graph 14:33–8, 27–8. doi:10.1016/0263-7855(96)00018-5

23. Jackson BM, Drysdale CM, Natarajan K, Hinnebusch AG. 1996. Identification of seven hydrophobic clusters in GCN4 making redundant contributions to transcriptional activation. Mol Cell Biol 16:5557–5571.

24. Jonker HRA, Wechselberger RW, Boelens R, Folkers GE, Kaptein R. 2005. Structural properties of the promiscuous VP16 activation domain. Biochemistry 44:827–839. doi:10.1021/bi0482912

25. Kabsch W, Sander C. 1983. Dictionary of protein secondary structure: pattern recognition of hydrogen-bonded and geometrical features. Biopolymers 22:2577–2637.

26. Kim J-Y, Chung HS. 2020. Disordered proteins follow diverse transition paths as they fold and bind to a partner. Science 368:1253–1257. doi:10.1126/science.aba3854

27. Kim J-Y, Meng F, Yoo J, Chung HS. 2018. Diffusion-limited association of disordered protein by non-native electrostatic interactions. Nat Commun 9:4707. doi:10.1038/s41467-018-06866-y

28. Kinney JB, Murugan A, Callan CG Jr, Cox EC. 2010. Using deep sequencing to characterize the biophysical mechanism of a transcriptional regulatory sequence. Proc Natl Acad Sci U S A 107:9158–9163. doi:10.1073/pnas.1004290107

29. Krois AS, Dyson HJ, Wright PE. 2018. Long-range regulation of p53 DNA binding by its intrinsically disordered N-terminal transactivation domain. Proc Natl Acad Sci U S A 115:E11302–E11310. doi:10.1073/pnas.1814051115

30. Krois AS, Ferreon JC, Martinez-Yamout MA, Dyson HJ, Wright PE. 2016. Recognition of the disordered p53 transactivation domain by the transcriptional adapter zinc finger domains of CREB-binding protein. Proc Natl Acad Sci U S A 113:E1853–62. doi:10.1073/pnas.1602487113

31. Kyte J, Doolittle RF. 1982. A simple method for displaying the hydropathic character of a protein. J Mol Biol 157:105–132.

32. Lambert SA, Jolma A, Campitelli LF, Das PK, Yin Y, Albu M, Chen X, Taipale J, Hughes TR, Weirauch MT. 2018. The Human Transcription Factors. Cell 175:598–599. doi:10.1016/j.cell.2018.09.045

33. Lando D, Peet DJ, Gorman JJ, Whelan DA, Whitelaw ML, Bruick RK. 2002. FIH-1 is an asparaginyl hydroxylase enzyme that regulates the transcriptional activity of hypoxia-inducible factor. Genes Dev 16:1466–1471. doi:10.1101/gad.991402

34. Latchman DS. 2008. Eukaryotic Transcription Factors, 5th Edition. ed. Elsevier Science.

35. Lecoq L, Raiola L, Chabot PR, Cyr N, Arseneault G, Legault P, Omichinski JG. 2017. Structural characterization of interactions between transactivation domain 1 of the p65 subunit of NF-κB and transcription regulatory factors. Nucleic Acids Res 45:5564–5576. doi:10.1093/nar/gkx146

36. Lin J, Chen J, Elenbaas B, Levine AJ. 1994. Several hydrophobic amino acids in the p53 amino-terminal domain are required for transcriptional activation, binding to mdm-2 and the adenovirus 5 E1B 55-kD protein. Genes Dev 8:1235–1246. doi:10.1101/gad.8.10.1235

37. Liu Y, Matthews KS, Bondos SE. 2008. Multiple intrinsically disordered sequences alter DNA binding by the homeodomain of the Drosophila hox protein ultrabithorax. J Biol Chem 283:20874–20887. doi:10.1074/jbc.M800375200

38. Ma J, Ptashne M. 1987. A new class of yeast transcriptional activators. Cell 51:113–119.

39. Maricque BB, Chaudhari HG, Cohen BA. 2018. A massively parallel reporter assay dissects the influence of chromatin structure on cis-regulatory activity. Nat Biotechnol. doi:10.1038/nbt.4285

40. Martin EW, Holehouse AS, Grace CR, Hughes A, Pappu RV, Mittag T. 2016. Sequence Determinants of the Conformational Properties of an Intrinsically Disordered Protein Prior to and upon Multisite Phosphorylation. J Am Chem Soc 138:15323–15335. doi:10.1021/jacs.6b10272

41. Metskas LA, Rhoades E. 2015. Conformation and Dynamics of the Troponin I C-Terminal Domain: Combining Single-Molecule and Computational Approaches for a Disordered Protein Region. J Am Chem Soc 137:11962–11969. doi:10.1021/jacs.5b04471

42. Oldfield CJ, Dunker AK. 2014. Intrinsically disordered proteins and intrinsically disordered protein regions. Annu Rev Biochem 83:553–584. doi:10.1146/annurev-biochem-072711-164947

43. Pace CN, Scholtz JM. 1998. A helix propensity scale based on experimental studies of peptides and proteins. Biophys J 75:422–427. doi:10.1016/s0006-3495(98)77529-0

44. Park M, Patel N, Keung AJ, Khalil AS. 2019. Engineering Epigenetic Regulation Using Synthetic Read-Write Modules. Cell. doi:10.1016/j.cell.2018.11.002

45. Piskacek S, Gregor M, Nemethova M, Grabner M, Kovarik P, Piskacek M. 2007. Nine-amino-acid transactivation domain: establishment and prediction utilities. Genomics 89:756–768. doi:10.1016/j.ygeno.2007.02.003

46. Powers SK, Holehouse AS, Korasick DA, Schreiber KH, Clark NM, Jing H, Emenecker R, Han S, Tycksen E, Hwang I, Sozzani R, Jez JM, Pappu RV, Strader LC. 2019. Nucleo-cytoplasmic Partitioning of ARF Proteins Controls Auxin Responses in Arabidopsis thaliana. Mol Cell 76:177–190.e5. doi:10.1016/j.molcel.2019.06.044

47. Raj N, Attardi LD. 2017. The Transactivation Domains of the p53 Protein. Cold Spring Harb Perspect Med 7. doi:10.1101/cshperspect.a026047

48. Ravarani CN, Erkina TY, De Baets G, Dudman DC, Erkine AM, Babu MM. 2018. High-throughput discovery of functional disordered regions: investigation of transactivation domains. Mol Syst Biol 14:e8190. doi:10.15252/msb.20188190

49. Regier JL, Shen F, Triezenberg SJ. 1993. Pattern of aromatic and hydrophobic amino acids critical for one of two subdomains of the VP16 transcriptional activator. Proc Natl Acad Sci U S A 90:883–887.

50. Rollins NJ, Brock KP, Poelwijk FJ, Stiffler MA, Gauthier NP, Sander C, Marks DS. 2019. Inferring protein 3D structure from deep mutation scans. Nat Genet 51:1170–1176. doi:10.1038/s41588-019-0432-9

51. Sanborn AL, Yeh BT, Feigerle JT, Hao CV, Townshend RJL, Aiden EL, Dror RO, Kornberg RD. 2020. Simple biochemical features underlie transcriptional activation domain diversity and dynamic, fuzzy binding to Mediator. Cold Spring Harbor Laboratory. doi:10.1101/2020.12.18.423551

52. Schmiedel JM, Lehner B. 2019. Determining protein structures using deep mutagenesis. Nat Genet 51:1177–1186. doi:10.1038/s41588-019-0431-x

53. Sharon E, Kalma Y, Sharp A, Raveh-Sadka T, Levo M, Zeevi D, Keren L, Yakhini Z, Weinberger A, Segal E. 2012. Inferring gene regulatory logic from high-throughput measurements of thousands of systematically designed promoters. Nat Biotechnol 30:521–530. doi:10.1038/nbt.2205

54. Sigler PB. 1988. Transcriptional activation. Acid blobs and negative noodles. Nature 333:210–212. doi:10.1038/333210a0

55. Staller MV, Holehouse AS, Swain-Lenz D, Das RK, Pappu RV, Cohen BA. 2018. A High-Throughput Mutational Scan of an Intrinsically Disordered Acidic Transcriptional Activation Domain. Cell Syst 6:444–455.e6. doi:10.1016/j.cels.2018.01.015

56. Tareen A, Kinney JB. 2020. Logomaker: beautiful sequence logos in Python. Bioinformatics 36:2272–2274. doi:10.1093/bioinformatics/btz921

57. Trojanowski J, Frank L, Rademacher A, Grigaitis P, Rippe K. 2021. Transcription activation is enhanced by multivalent interactions independent of liquid-liquid phase separation. doi:10.1101/2021.01.27.428421

58. Tycko J, DelRosso N, Hess GT, Aradhana, Banerjee A, Mukund A, Van MV, Ego BK, Yao D, Spees K, Suzuki P, Marinov GK, Kundaje A, Bassik MC, Bintu L. 2020. High-Throughput Discovery and Characterization of Human Transcriptional Effectors. Cell 183:2020–2035.e16. doi:10.1016/j.cell.2020.11.024

59. Vitalis A, Pappu RV. 2009. ABSINTH: a new continuum solvation model for simulations of polypeptides in aqueous solutions. J Comput Chem 30:673–699. doi:10.1002/jcc.21005

60. Warfield L, Tuttle LM, Pacheco D, Klevit RE, Hahn S. 2014. A sequence-specific transcription activator motif and powerful synthetic variants that bind Mediator using a fuzzy protein interface. Proc Natl Acad Sci U S A 111:E3506–13. doi:10.1073/pnas.1412088111

61. Wojciak JM, Martinez-Yamout MA, Dyson HJ, Wright PE. 2009. Structural basis for recruitment of CBP/p300 coactivators by STAT1 and STAT2 transactivation domains. EMBO J 28:948–958. doi:10.1038/emboj.2009.30

62. Yasinska IM, Sumbayev VV. 2003. S-nitrosation of Cys-800 of HIF-1α protein activates its interaction with p300 and stimulates its transcriptional activity. FEBS Lett 549:105–109.

